# Restoration ecologists might not get what they want: Global change shifts trade-offs among ecosystem functions

**DOI:** 10.1101/2020.12.21.423790

**Authors:** Sebastian Fiedler, José A.F. Monteiro, Kristin B. Hulvey, Rachel J. Standish, Michael P. Perring, Britta Tietjen

**Affiliations:** Freie Universität Berlin, Theoretical Ecology, Institute of Biology, Königin-Luise-Straße 2/4, Gartenhaus, 14195 Berlin, Germany; Berlin-Brandenburg Institute of Advanced Biodiversity Research (BBIB), 14195 Berlin, Germany; University Bayreuth, Department of Ecological Modelling, 95448 Bayreuth, Germany; Statistical Office Basel-Stadt, Basel, Switzerland; Working Lands Conservation, 1460 E 1220, Logan, UT 84341, USA; Environmental and Conservation Sciences, Murdoch University, 90 South Street, Murdoch, WA 6150, Australia; Forest & Nature Lab, Ghent University, BE-9090, Gontrode-Melle, Belgium; Ecosystem Restoration and Intervention Ecology Research Group, School of Biological Sciences, The University of Western Australia, 35 Stirling Highway, Crawley, WA 6009, Australia; Centre for Ecology and Hydrology, Environment Centre Wales, Deiniol Road, Bangor, Gwynedd, LL57 2UW, United Kingdom

**Keywords:** Biodiversity, Functional Traits, Plant Traits, Climate Change, Ecosystem Services, Mediterranean-type Ecosystem, Multifunctionality, Simulation Model

## Abstract

1. Ecological restoration increasingly aims at improving ecosystem multifunctionality and making landscapes resilient to future threats, especially in biodiversity hotspots such as Mediterranean-type ecosystems. Successful realisation of such a strategy requires a fundamental mechanistic understanding of the link between ecosystem plant composition, plant traits and related ecosystem functions and services, as well as how climate change affects these relationships. An integrated approach of empirical research and simulation modelling with focus on plant traits can allow this understanding.
2. Based on empirical data from a large-scale restoration project in a Mediterranean-type climate in Western Australia, we developed and validated the spatially explicit simulation model ModEST, which calculates coupled dynamics of nutrients, water and individual plants characterised by traits. We then simulated all possible combinations of eight plant species with different levels of diversity to assess the role of plant diversity and traits on multifunctionality, the provision of six ecosystem functions (covering three ecosystem services), as well as trade-offs and synergies among the functions under current and future climatic conditions.
3. Our results show that multifunctionality cannot fully be achieved because of trade-offs among functions that are attributable to sets of traits that affect functions differently. Our measure of multifunctionality was increased by higher levels of planted species richness under current, but not future climatic conditions. In contrast, single functions were differently impacted by increased plant diversity. In addition, we found that trade-offs and synergies among functions shifted with climate change.
4. *Synthesis and application*. Our results imply that restoration ecologists will face a clear challenge to achieve their targets with respect to multifunctionality not only under current conditions, but also in the long-term. However, once ModEST is parameterized and validated for a specific restoration site, managers can assess which target goals can be achieved given the set of available plant species and site-specific conditions. It can also highlight which species combinations can best achieve long-term improved multifunctionality due to their trait diversity.

## 1 INTRODUCTION

Global change is contributing to a decline in biodiversity, and ecosystem functions and services people rely on for well-being (IPBES, 2019). Degradation associated with past change, and concern for the future supply of multiple ecosystem services is particularly apparent in Mediterranean-type ecosystems where remarkably high diversity is threatened by multiple environmental changes (Cowling et al., 1996; Sala, 2000). Ecological restoration is therefore required in such systems, with the goal to maintain the long-term and simultaneous delivery of multiple ecosystem functions and services in these socio-ecological systems (Shackelford et al., 2013; Gann et al., 2019).

Managing landscapes for multiple functions or services simultaneously requires a direct comparison of their delivery (e.g. Byrnes et al., 2014; Manning et al., 2018). With increasing evidence that higher levels of ecosystem functions and services are associated with greater species numbers (Cardinale et al., 2012), the traditional focus of restoration on plant biodiversity appears justified (Perring et al., 2015). Enhanced biodiversity, however, does not necessarily increase the simultaneous and resilient provision of multiple ecosystem services as some services do not profit from greater species richness and the effect of global change on species and service provisioning remains unclear (e.g. Cardinale et al., 2012; Meyer et al., 2018).

In an attempt to further understanding of biodiversity’s impact on ecosystems, restoration ecology has more recently made use of the functional trait concept allowing selection of plant species based on their response and effect traits (Lavorel & Garnier, 2002; Laughlin, 2014). A focus on effect traits, which have been found to be linked to ecosystem functions (Lavorel & Garnier, 2002), allows for a better comparison across individuals and plant species. We have a good understanding on how individual environmental factors affect individual functions and services via plant traits (e.g. Lavorel & Garnier, 2002; Suding et al., 2008). However, nature is more complex and plant traits are not always linked to single functions. Instead, multiple traits can affect one function, and multiple functions can be affected by a single trait (de Bello et al., 2010), and multiple functions can influence a single ecosystem service (Fu et al., 2013). This is particularly important if traits positively affect one function while at the same time negatively impacting another one – so-called trade-offs (Bennett et al., 2009). Knowing the trade-offs as well as synergies among plant traits and functions is therefore important for selecting plant species based on their traits to simultaneously improve levels of multiple functions/services.

In addition, multiple environmental change factors that directly, or indirectly (via altering plant trait distributions), affect ecosystem functions can have non-additive effects (e.g. Luo et al. 2008). Restoration strategies based on individually studied effects could therefore be problematic when trying to achieve a long-term supply of functions and services. Furthermore, traits within a plant community may be affected differently by environmental factors, and therefore the provision of trait-mediated ecosystem functions may be affected differently as well. Consequently, trade-offs among ecosystem services observed under current environmental conditions might not be the same under future conditions.

To improve understanding and allow more informed restoration, Fiedler et al. (2018) suggested an integrated approach that focuses on plant traits and combines the strengths of empirical and simulation modelling studies. Empirical approaches can support modelling approaches with essential data, while simulation models can extend empirical approaches by allowing assessment of the multi-layered relationship between multiple environmental factors, plant traits and ecosystem functions/services on larger temporal and spatial scales. Current trait-based simulation models provide a good basis for this approach (e.g. Esther et al., 2011; Fyllas & Troumbis, 2009; Schaphoff et al., 2017). However, to be able to support restoration towards multifunctional and resilient ecosystems, simulation models need to be combined and extended to meet the following criteria: (i) coupled processes for soil water, nutrient and plants as well as the respective feedbacks allowing to mechanistically study the impact of global change on vegetation, (ii) consideration of individual interactions (e.g. facilitation and competition) as well as spatial heterogeneity relevant for applied restoration projects implemented on smaller spatial scales, and (iii) a thorough validation of model outcomes against field data to make simulation models applicable for restoration.

Based on existing model tools and a restoration experiment in a Mediterranean-type ecosystem in SW Australia (Perring et al., 2012), we therefore developed and validated the individual- and trait-based simulation model ModEST (Modelling Ecosystem Functions and Services based on Traits). ModEST links water, nitrogen and plant processes dependent on climatic and other environmental conditions and exhibits therefore enough generality to transfer findings beyond this specific study site. We used the model to assess the following research questions:

1. What is the role of planted species richness under current and future conditions on multifunctionality, and the provision of separate ecosystem functions and services?
2. How will environmental changes affect trade-offs and synergies among ecosystem functions and services of simulated plant communities?
3. What sets of plant traits and correlations among them in the simulated plant communities provide ecosystem functions under current and future conditions?

## 2 MATERIAL AND METHODS

### 2.1 Model description

We developed a spatially explicit model, ModEST (**Mod**elling **E**cosystem Functions and **S**ervices based on **T**raits) which simulates the coupled daily dynamics of nutrients, water, and individual woody plants (Fig. 1), from which different ecosystem functions and services can be estimated (Fiedler et al., 2020). The model landscape is subdivided into grid cells (5 × 5 m^2^), two soil layers, and individual plants characterized by coordinates within the landscape. The model runs for different environmental settings concerning soil texture, climatic conditions, topography, initial plant composition and their traits, with full descriptions given in Supplementary S1 and S2. In the following, we briefly describe the three coupled modules of ModEST.

**Figure 1:**
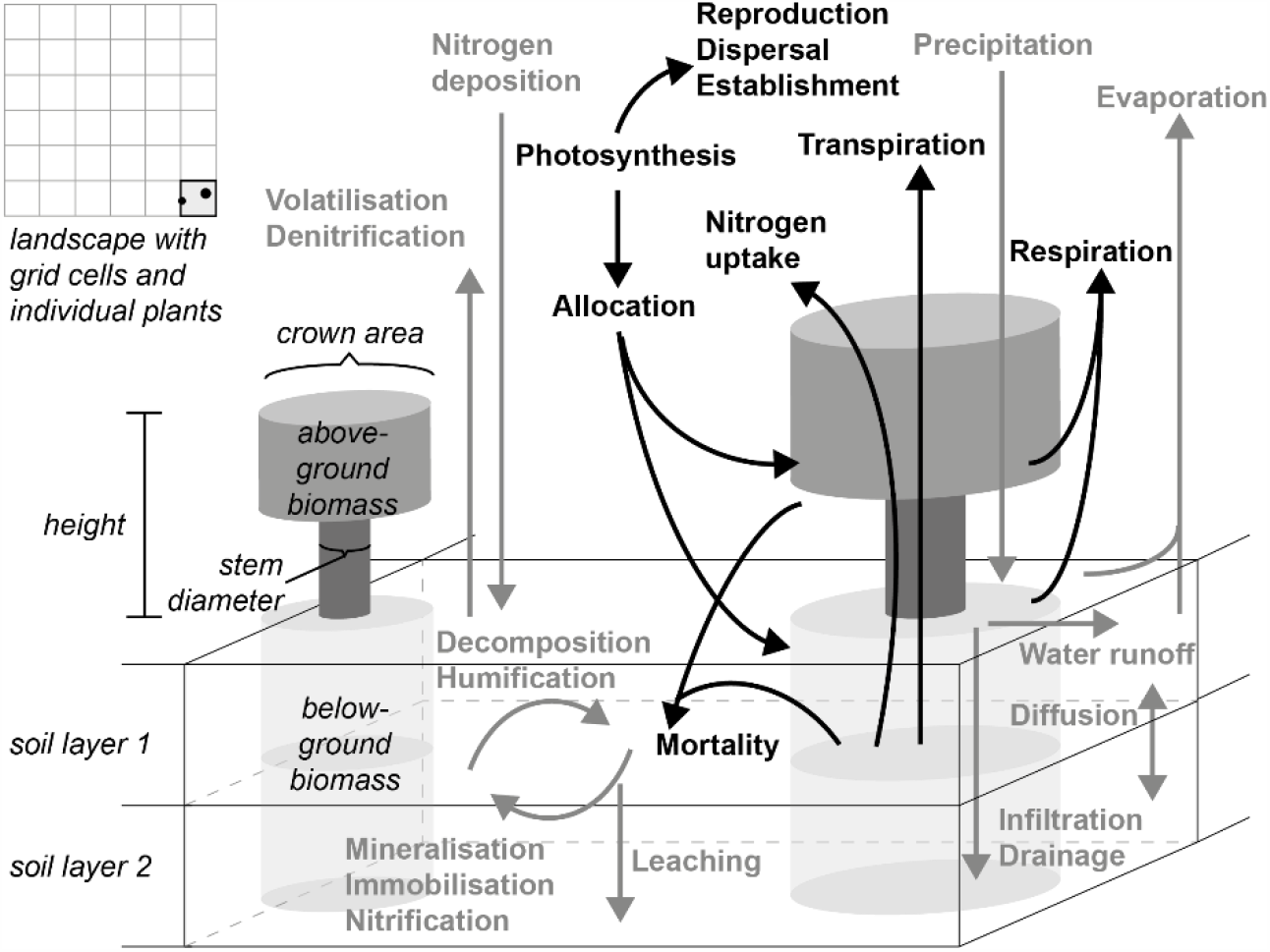
Structure (*italic*) and processes (bold) of ModEST. The modelled landscape is sub-divided into grid cells consisting of two soil layers as well as individual woody plants that are characterized by above- and below-ground features and are continuously distributed over the landscape. Coupled processes are calculated, i.e. hydrological and nutrient processes for each grid cell and soil layer (bold grey) as well as plant processes for each individual plant (bold black) depending on the resources of its covering grid cell.

The nutrient module is based on processes for simulating soil nitrogen and soil carbon described in the model SWAT (Kemanian et al., 2011). Daily dynamics of soil organic matter (SOM), nitrate, and ammonium in two soil layers are driven by *nitrogen deposition* from the atmosphere, *decomposition* and *humification* of plants’ residue to SOM, *immobilization, mineralization* to ammonium, *nitrification* to nitrate as well as nutrient losses through *volatilization, denitrification* and *leaching*.

We based the hydrological module on the approach of Tietjen et al. (2009), who simulated surface water and soil moisture in two soil layers. Daily water dynamics are driven by *precipitation, lateral water redistribution of surface water, infiltration*, and *vertical fluxes*, and by water losses via *evaporation* and *transpiration*. For ModEST, we adopted these processes with the exception of transpiration which we implemented after the model LPJ (Sitch et. al., 2003) and LPJmL (Schapoff et al., 2017), to better account for stomatal conductance (see description of the transpiration process in Supplementary S1).

The plant module is mainly based on LPJ and LPJmL (Schaphoff et al., 2017; Sitch et al., 2003; Smith et al., 2014) and local processes as described for an individual-based plant model by May et al. (2009). The module simulates the life cycle of individual woody plants placed in the landscape, their dynamic below- and aboveground carbon and nitrogen pools as well as structural components (e.g. plant height, crown area) based on plant traits and abiotic conditions. We adopted – with some changes – the plant processes *photosynthesis, transpiration, respiration, reproduction*, and *allocation* after Sitch et. al. (2003) and Schapoff et al. (2017), *nitrogen uptake* after Smith et al. (2014), as well as *dispersal and establishment* after May et al. (2009). We added a simple *plant mortality* process based on annual plant growth and a species-specific growth threshold below which the individual plant dies. Given these adaptations, we fully describe this module in the Supplementary S1.

### 2.2 Model parameterization and validation

We parameterised and validated ModEST based on the settings of the Ridgefield experiment, a large-scale restoration experiment situated in the wheatbelt of SW Australia on former agricultural land (Perring et al., 2012). The experiment is located in a Mediterranean-climate region (32°29’S 116°58’E, elevation 350 m a.s.l.) with mean annual rainfall of 453 mm (2013 – 2019) and precipitation mainly during winter. The average maximum daily temperature in January is 30.7 °C and the average minimum daily temperature in August is 7.6 °C.

We parameterized eight evergreen woody plant species (Table S2.1) which were planted in different plant assemblages representing increasing functional and species richness. We used the most prevalent soil type (loamy sand, Table S2.2) in the experiment (see Supplementary S2 for full description of model parameterisation).

For model validation, we checked the outcome of the parameterized model against measurements from Ridgefield plots (see Supplementary S2 for model settings). We quantitatively compared simulated and observed dynamics using Spearman’s rank correlation *r* and the root mean square error *RMSE*. Simulated aboveground alive biomass, mean plant height, and surviving individual counts agreed well with the measured data (i.e. significant [p < 0.01] correlations, low RMSE). Exceptions were the biomass dynamic of *B. sessilis* and the population dynamics of *C. quadrifidus* and *C. phoeniceus*, where correlations were insignificant (Fig. S2.2). However, RMSE for these cases remained low (RMSE < 1.0), indicating only small deviances between simulated and measured dynamics, and suggesting reasonable model behavior.

### 2.3 Simulation experiments

We simulated a full-factorial design of plant species combinations using the eight species included in the Ridgefield study (and thus simulating plant assemblages beyond those planted at Ridgefield) to assess ecosystem functioning under current and future climatic conditions. The flat modelled landscape (50 × 50 m^2^) contained a homogenous soil texture of loamy sand, with initial soil moisture (= 0.15 m^3^ · m^-3^), ammonium (= 2.35 mg · kg^-1^) and nitrate (= 9.92 mg · kg^-1^) set to the mean measured values across all Ridgefield plots with soil texture loamy sand. Each scenario was repeated ten times to account for stochasticity in the initialisation of plant individuals (see *Species richness scenarios*), weather input (see *Climate change scenarios*), and the dispersal process (see model description in Supplementary S1).

#### 2.3.1 Species richness scenarios

All possible combinations of the eight woody plant species used in the Ridgefield experiment were simulated leading to 255 different plant species compositions. Using this design, communities covered a wide range of different plant trait combinations, and species richness varied from monocultures to 8-species mixtures. For each simulation, 500 one-year old individuals with the same or a similar initial individual number of each present species were randomly positioned with 2 m distance to neighbouring individuals. Initial plant heights were randomly drawn from a species-specific normal distribution that was obtained from height distributions of the one-year planted individuals in the Ridgefield experiment (Fig. S3.1).

#### 2.3.2 Climate change scenarios

For current climatic conditions, we used corrected daily precipitation, minimum and maximum air temperature and solar radiation data from 1990 to 2018 from the weather station in Pingelly (32°31’S 117°04’E, 297 m a.s.l.) about 12 km away from our study site (Bureau of Meteorology, 2019, Supplementary S3.1). Atmospheric CO_2_ was set to 400 ppm.

For assessing impacts of climate change, we obtained the anomalies for future conditions (2080 – 2099) compared to past conditions (1986 – 2005) separately for each season based on the four climate projection Representative Concentration Pathways (RCPs) for SW Australia (Hope et al., 2015). We added the median reported trend between past and future climate from different global climate model simulations to the current weather data from Pingelly to generate realistic time series of future weather data. Atmospheric CO_2_ was set according to IPCC (2014).

For each model repetition, we randomly selected yearly weather data from the current or future weather data set, given the climate scenario, to get 50 years of weather time-series input data.

As we found qualitatively similar patterns on ecosystem functioning across the different RCPs (Fig. S4.1), we focused, for better clarity, on the most extreme climate projection RCP 8.5 with an increase in mean annual air temperature by 3.4 °C and a decrease in mean annual precipitation by 16 % (Table S3.1, Fig. S3.2).

#### 2.3.3 Evaluation of simulation outcomes

To assess the provision of, and trade-offs and synergies among, ecosystem functions, we determined the supply of six functions chosen to cover three major ecosystem services, namely *water supply, nutrient supply*, and *carbon sequestration* (Table 1). *Water supply* involves two functions: groundwater recharge – approximated by deep drainage of water below 2 meters soil depth – and ecosystem water use efficiency – the net primary productivity of the ecosystem per unit precipitation. *Nutrient supply* is represented by ecosystem nitrogen use efficiency – estimated by the net primary productivity of the ecosystem per soil available nitrogen – and by litter quality – approximated by the nitrogen to carbon ratio (N:C) of the plant’s residue in the ecosystem originating from plant senescence and mortality. *Carbon sequestration* covers both plant and soil carbon increments.

**Table 1:** Ecosystem functions.

| Related ecosystem service | Ecosystem function | Model output | Unit |
| --- | --- | --- | --- |
| Water supply | Groundwater recharge (GWR) | Annual deep (> 2 meters in soil depth) soil water drainage per m<sup>2</sup> | mm · year<sup>-1</sup> |
|  | Ecosystem water use efficiency (EWU) | Annual net primary productivity (NPP) per m<sup>2</sup>/ Annual precipitation per m<sup>2</sup> | g · L<sup>-1</sup> · year<sup>-1</sup> |
| Nutrient supply | Ecosystem nitrogen use efficiency (ENU) | Annual NPP per m<sup>2</sup>/ Annual mean soil avail. nitrogen per m<sup>3</sup> | kgNPP · m<sup>-2</sup> · kgN<sup>-1</sup> · m<sup>-3</sup> |
|  | Ecosystem litter quality (ELQ) | Annual nitrogen per m<sup>2</sup>/ Annual carbon per m<sup>2</sup> | gN · year<sup>-1</sup> · kgC<sup>-1</sup> · year<sup>-1</sup> |
| Carbon sequestration | Total plant carbon increment (PCI) | Annual plant carbon increment | kg · m<sup>-2</sup> · year<sup>-1</sup> |
|  | Total soil carbon increment (SCI) | Annual soil carbon increment | t · m<sup>-2</sup> · year<sup>-1</sup> |

Across these six functions we estimated the multifunctionality as suggested by van der Plas et al. (2016). Accordingly, all ecosystem functions were standardized between 0 and 1 based on the minimum and maximum value per function in a given climate scenario as well as across climate scenarios. Multifunctionality was then defined as the number of functions above a threshold for each species richness and climate change scenario. We chose a threshold level of 50 % which has been shown to be comparable across different countries (van der Plas et al., 2016).

We evaluated plant trait distribution through calculating community weighted mean (CWM) for selected traits (Table 2). These traits are measurable in the field and therefore applicable for ecosystem restoration.

**Table 2:**
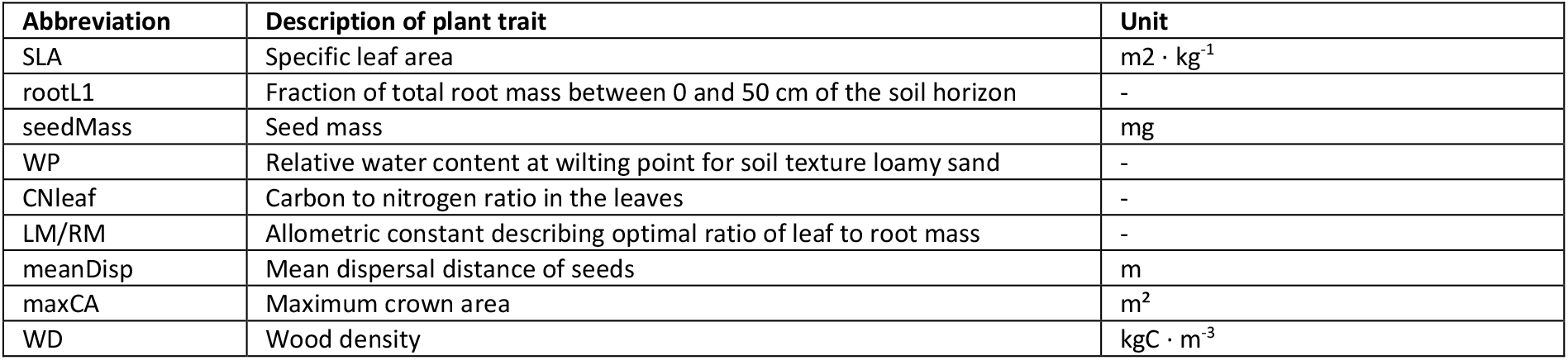
Focal plant traits.

| Abbreviation | Description of plant trait | Unit |
| --- | --- | --- |
| SLA | Specific leaf area | m<sup>2</sup> · kg<sup>-1</sup> |
| rootL1 | Fraction of total root mass between 0 and 50 cm of the soil horizon | - |
| seedMass | Seed mass | mg |
| WP | Relative water content at wilting point for soil texture loamy sand | - |
| CNleaf | Carbon to nitrogen ratio in the leaves | - |
| LM/RM | Allometric constant describing optimal ratio of leaf to root mass | - |
| meanDisp | Mean dispersal distance of seeds | m |
| maxCA | Maximum crown area | m<sup>2</sup> |
| WD | Wood density | kgC · m<sup>-3</sup> |

We evaluated model outcomes between 40 and 50 years given attainment of dynamic equilibrium in total plant species cover after 40 years (Fig. S3.3). All relationships were analysed by a Spearman’s rank correlation.

## 3 RESULTS

### 3.1 Planted species richness effects on ecosystem functioning

With higher planted and realised richness, we found that our approximation of ecosystem multifunctionality increased under current climatic conditions but decreased under future conditions (Fig. 2A, left; see also Fig. S4.2). However, when considering current and future knowledge for the standardisation of ecosystem functions, current and future multifunctionality decreased with greater richness (Fig. 2A, right).

**Figure 2:**
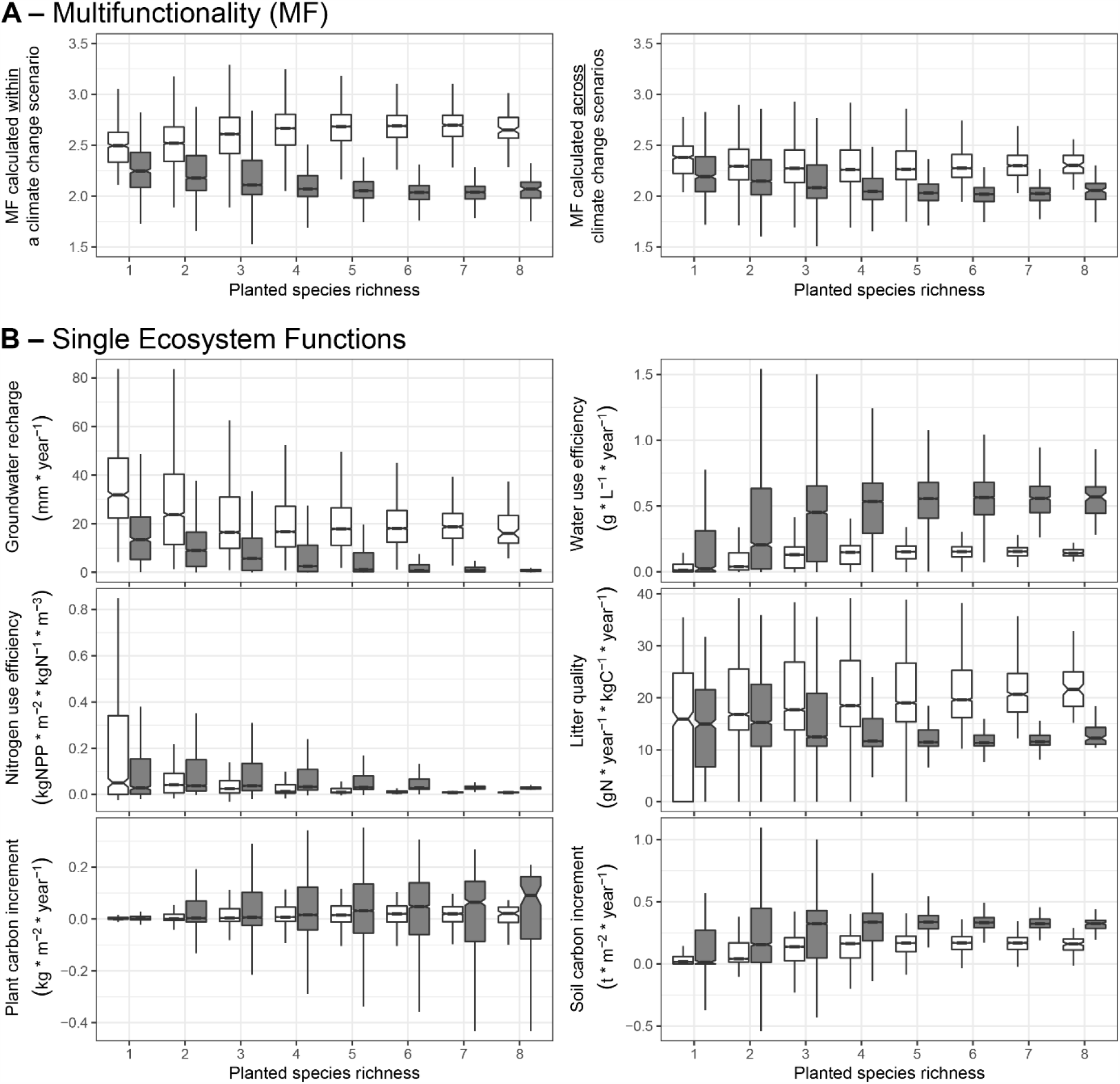
Multifunctionality (A) and single ecosystem functioning (B) for each planted species richness under current (white boxplots) and future climatic conditions (grey boxplots). Multifunctionality is either calculated within each climate scenario (A, left) or across climate scenarios (A, right). Shown is functioning for the last 10 simulated years and for 10 model repetitions as well as for 255 different plant communities which are unevenly distributed across the different planted species richness scenarios according to maximal possible combinations out of the pool of eight focal plant species. For better comparability among boxplots, single outliers are not shown.

Some single ecosystem functions increased but others decreased with planted species richness under current conditions (Fig. 2B). Climate change strengthened this pattern and increased variability for most of the functions, except for groundwater recharge and litter quality. For communities with up to three or four planted species, groundwater recharge decreased, whereas the water use efficiency of the ecosystem increased. If more than three or four species were planted, both functions remained stable. Nitrogen use was most efficient for monocultures. In contrast, litter quality increased with higher planted richness under current conditions reaching maximum quality for the most speciose community, while under future conditions litter quality decreased with higher planted richness. Soil carbon, and to a lesser extent plant carbon, increments, were enhanced with higher planted richness, reaching their maximum at an intermediate richness, and remaining stable for higher values. Except for plant carbon increment, all ecosystem functions showed a decreasing variability with increasing planted richness (Figs 2B and S4.3).

### 3.2 Trade-offs and synergies among ecosystem functions

With the eight plant species considered in this study, ecosystem multifunctionality could not fully be achieved, in current or future conditions (MF much smaller than 6, Fig. 2A), since there are negative correlations (trade-offs) among functions (Fig. 3). Multifunctionality benefited from a strong positive correlation (synergy) between soil carbon increment and water use (Figs 2B and 3). However, stronger trade-offs between ecosystem nitrogen use and litter quality as well as between groundwater recharge and ecosystem water use or soil carbon increment mostly constrained the maximisation of the multifunctionality.

**Figure 3:**
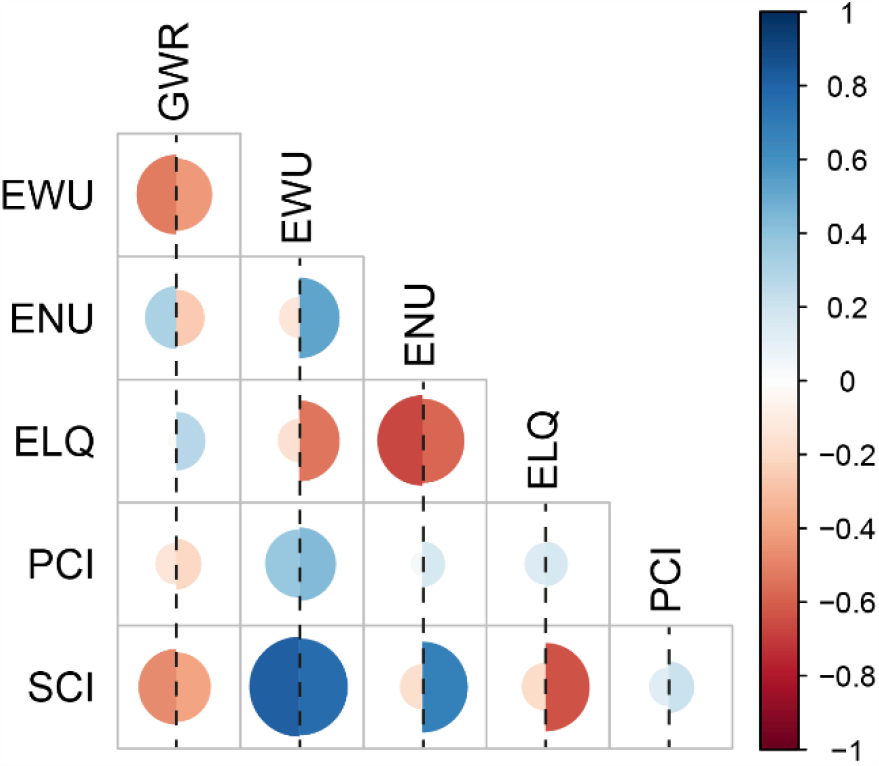
Negative (trade-off, red) and positive (synergy, blue) relationships among ecosystem functions under current (left half circle) and future climatic conditions (right half circle). Shown are significant Spearman’s rank correlations (α = 0.05) among ecosystem functions based on the last 10 simulated years and for 10 model repetitions across all 255 simulated plant communities. Meaning of abbreviations can be found in Table 1.

Most relationships between nitrogen use efficiency and other functions reversed under future conditions: in contrast to current conditions, an increase in nitrogen use efficiency was now accompanied by a decline in groundwater recharge as well as a strong increase in water use and soil carbon increment in the ecosystem. In addition, ecosystem litter quality and groundwater recharge could be increased at the same time under future conditions, which was not possible under current conditions. Some trade-offs and synergies observed under current conditions did not reverse but rather strengthened under the future climate scenario: trade-offs between ecosystem litter quality and ecosystem water usage, or soil carbon increment, became more apparent, whereas ecosystem nitrogen use efficiency and plant carbon increment were more effectively maximised at the same time.

### 3.3 Plant traits in the community and ecosystem functioning

Community weighted mean plant traits could be linked to single ecosystem functions (Fig. 4A). Particular trait combinations rather than single traits affected individual functions. Functions within the ecosystem services of water and nutrient supply showed contrasting correlations to plant traits in the community, explaining their strong trade-offs. For example, under current conditions groundwater recharge (GWR) was enhanced by communities with a low specific leaf area (SLA), higher investment into leaves than into roots (LM/RM), smaller crowns (maxCA), lower wood density (WD), and a higher wilting point (WP). In contrast, to achieve a maximal ecosystem water use efficiency (EWU), wood density and maximum crown area should be larger coupled with a deeper rooting system (low value of rootL1). Almost the same features that maximised ecosystem water use efficiency also increased plant carbon increment (PCI) and soil carbon increment (SCI) in the ecosystem, supporting the synergies among the three functions.

**Figure 4:**
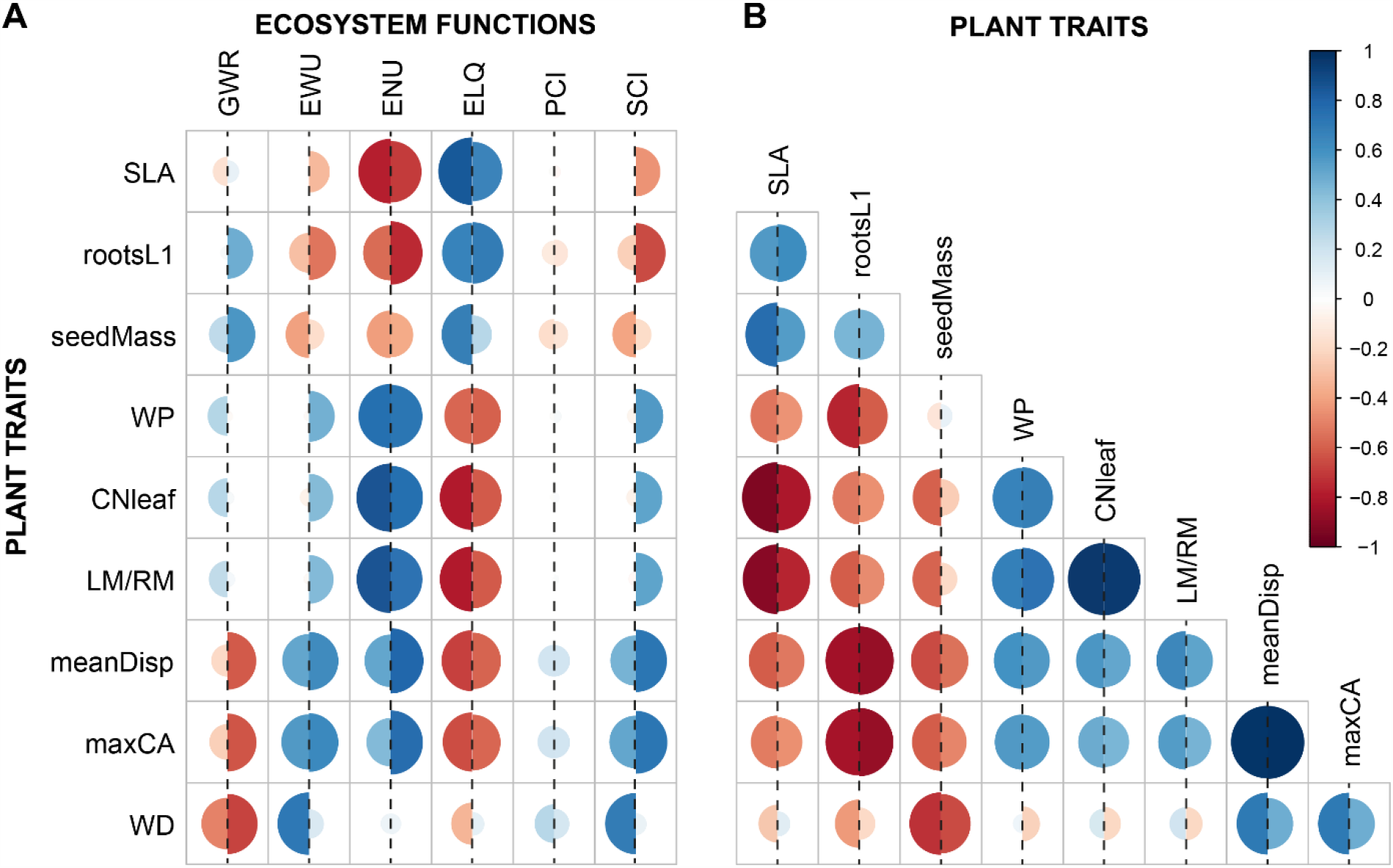
Plant trait – ecosystem function relationships and intrinsic community trait correlations under current (left half circle) and future climatic conditions (right half circle). Spearman’s rank correlations (A) between CWM plant traits and ecosystem functions, and (B) among CWM plant traits. Shown are significant correlations (α = 0.05) based on the last 10 simulated years and for 10 model repetitions across all 255 simulated plant communities. Meaning of abbreviations can be found in Table 1 for ecosystem functions and Table 2 for CWM plant traits.

Under future climatic conditions, traits gained or lost their importance especially for soil carbon increment and functions related to water supply. Other functions, i.e. nitrogen use efficiency, traits associated with ecosystem litter quality and plant carbon increment showed no or limited change in importance. For example, under future conditions, wilting point, carbon to nitrogen ratio in leaves, and leaf to root mass ratio gained importance for restoring water use efficiency. The same traits remained important for restoring nitrogen use efficiency, which might explain the shift from a trade-off to a synergy between these two functions. However, most trait-trait correlations remained the same except for correlations involving wood density and wilting point (Fig. 4B) suggesting that changes in the relationships among functions was not driven by underlying changes in trait-trait correlations. Still, we found that trait compositions shifted with climate change in particular for more speciose communities (Figs S4.4 and S4.5), i.e. shifts to plants with deeper roots, higher maximal crown area and with lighter and far-dispersed seeds. These changes led to a stronger decrease in groundwater recharge and ecosystem litter quality (Figs 2B and 4A), which explains the decreasing multifunctionality with increasing planted richness under climate change (Fig. 2A).

## 4 DISCUSSION

### 4.1 Trade-offs prevent maximised multifunctionality

We found that trade-offs prevented the achievement of restoration goals with simultaneous maximisation of multiple functions and services. Our integrated empirical-modelling approach allowed us to reveal the mechanisms. Trade-offs emerged for two reasons: the same trait or group of traits can have positive effects on one function, but negative effects on a second function (e.g. de Bello et al., 2010; Teixeira et al., 2020), and the correlation of plant traits can mean that some traits reliably coexist, while some never do (e.g. de la Riva et al., 2016).

Even though we could not achieve full multifunctionality in our study, i.e. all functions at their maximum, we found that in a restoration setting, bundles of functions with synergies among them could be maximised instead. For instance, if managers want to improve ecosystem water use efficiency and carbon sequestration (but not nitrogen supply), this can be achieved by planting communities with deeper roots, greater crown area and wood density as well as small seeds with larger dispersal distances. However, the two functions defining the service ‘water supply’ in our study, i.e. ecosystem water use efficiency and groundwater recharge, could not be optimized at the same time. Thus, the choice of the ecosystem services to be restored might be very crucial. Since only bundles of services can be optimized at the same time, different bundles could be integrated across the landscape to achieve landscape multifunctionality (e.g. Lovell & Johnston, 2009).

### 4.2 Trade-offs among functions shift with climate change

We found that trade-offs and synergies among ecosystem functions observed under current conditions shifted under future conditions, posing a clear challenge for long-term restoration outcomes. These shifts in the relationships among functions can be explained either by a direct change of ecosystem functioning differently affected by changing environmental conditions or indirectly through uneven shifts in underlying community plant traits and thus changes in the correlations among CWM traits (e.g. Zirbel et al., 2017). In this context, we should note that the model did not incorporate trait plasticity, which might attenuate or enhance shifts in relationships. In our study, simulated climate change altered species and thus trait compositions as reviewed also by Maestre et al. (2012a) for drylands as well as single trait-trait correlations as also shown by Ahrens et al. (2020). However, most correlations among the traits within the community remained as observed under current conditions. Therefore, shifts in the relationships among functions were mostly due to a direct climate change impact, such as the simulated decrease in groundwater recharge via less available water for infiltration, and higher evapotranspiration due to warmer temperatures (cp. Reinecke et al., 2020) and a simultaneous increase in ecosystem nitrogen use efficiency via a carbon fertilisation effect (cp. Leakey et al., 2009), leading to the observed trade-off among the two functions under future climate.

### 4.3 Multifunctionality might not always be the right choice

If restoration aims to only increase but not maximise ecosystem multifunctionality, we found that promoting plant diversity achieved this goal, at least for our measure of multifunctionality and under current climatic condition. This is in line with previous findings and different measures of multifunctionality (Gross et al., 2017; Maestre et al., 2012b). However, a comparison of our chosen measure of multifunctionality with two other measures showed that our findings are not always apparent (Fig. S4.6, see also review by Maestre et al., 2012b and Manning et al., 2018). In addition, we found that the environmental context can significantly affect multifunctionality outcomes. For instance, in our study, minimum and maximum functioning, needed for the standardisation of each function, differed within or across climate scenarios and thus resulted into different patterns depending on the scope we looked at.

Furthermore, even though current multifunctionality in our study was improved by greater richness, single functions such as ecosystem nitrogen use efficiency were not. This specific finding contrasts with an empirical study that has shown complementary effects of diverse woody plant communities (mixtures of up to 16 evergreen and deciduous plant species) on nitrogen use (Schwarz et al., 2014). Here, we focused on eight evergreen woody species with similar C:N ratios (Table S2.1), thus complementary nitrogen use was likely not prevalent.

Greater planted richness decreased variability in ecosystem functioning suggesting a more consistent supply across the species combinations planted. This could be due to functional redundancy acting as stabilizing effect for a resilient supply of ecosystem functions (Mori et al., 2013). Under future conditions, however, higher plant diversity did not show greater resilience to environmental changes. Instead, we observed that with climate change speciose communities experienced greater species losses, potentially through higher interspecific competition (Ruiz-Benito et al., 2013), which in turn significantly lowered functional redundancy and thus the potential higher resilience against environmental changes. Also, even though multifunctionality decreased with higher planted richness under future conditions, only single functions, i.e. ecosystem litter quality, were largely affected and contributed to this decline, whereas most of the other functions increased with richness. Thus, the choice of metrics for restoration success should be considered if the goal is to improve a set of equally desired ecosystem functions and services at the same time.

### 4.4 Broader applicability of this study for restoration world-wide

We simulated the long-term effect of plant choice on multifunctionality and six separate ecosystem functions, grouped to the ecosystem services water supply, nutrient supply, and carbon sequestration, in a Mediterranean site in SW Australia. We found that the ultimate aim to improve restoration outcomes with respect to maximizing multiple ecosystem functions and services at the same time under current and future climatic conditions was limited by trade-offs among ecosystem functions which shifted with climate change.

Even though we focused on a specific Mediterranean site, we believe that our general interpretations pertain to terrestrial systems globally since underlying mechanisms driving trade-offs among functions and shifts in the trade-offs have been fundamentally shown across different ecosystems: i.e. ecosystem functions are affected by underlying plant traits (e.g. de Bello et al., 2010; Funk et al., 2017) and environmental change either directly or indirectly, via changing plant trait compositions, affects ecosystem functions (e.g. De Deyn et al., 2008; Garnier et al., 2007). Thus, restoration ecologists across the world will face a clear challenge to achieve their targets under current conditions and in the long-term.

However, ecosystem functioning as well as trade-offs among functions and how these will shift with climate change is likely to be context-dependent (e.g. dependent on local species pool, soil texture, weather, and regional projected climate change) and thus different across ecosystems (e.g. Ding et al., 2020; Ratcliffe et al., 2017). In addition, various ecosystems are degraded differently, and thus restoration managers might want to improve different desired functions.

With this study we applied the steps suggested by Fiedler et al. (2018) in order to improve ecological restoration and showed that models like ModEST can serve as a planning tool to better understand the suite of desired ecosystem functions and services that can be restored in any particular place based on the plant species available and the local environmental conditions. If further developed and combined with a graphical user interface and special training, this can help managers to set realistic restoration goals and select from a list of potential species and traits to achieve these goals.

## 5 AUTHORS’ CONTRIBUTION

SF, MP and BT conceived the project. SF implemented the hydrological and plant module, parameterized ModEST, conducted the experiments, analyzed the simulation outcomes, prepared all figures and tables, and wrote the first draft of the manuscript in close collaboration with BT and MP. JM implemented the nutrient module. KH, MP and RS participated in the conception and implementation of the Ridgefield experiment. All authors interpreted data, contributed critically to drafts, and gave final approval for publication.

## 6 ACKNOWLEDGEMENTS

This work was supported by the German Research Foundation (DFG project TI 824/3-1), the German Academic Exchange Service, and the University Alliance for Sustainability. The set-up of Ridgefield was supported by an Australian Research Council Laureate Fellowship to Richard J. Hobbs and funding from The University of Western Australia; UWA and the ARC Centre of Excellence for Environmental Decisions provide continued support. MPP was supported by the European Research Council PASTFORWARD project, awarded to Kris Verheyen (ERC Consolidator Grant 614839). The authors would like to thank the HPC Service of ZEDAT at Freie Universität Berlin for computing time, Marie-Sophie Rohwäder and Jonas Roth for data processing, Florian Hartig for his support on the Bayesian R package, Selina Baldauf, Katja Irob and Felix Wesener for their feedback on the manuscript as well as Rebecca Campbell and Tim Morald for their support regarding the Ridgefield experiment.

## 7 DATA AVAILABILITY STATEMENT

The source code of ModEST available from GitHub https://doi.org/10.5281/ZENODO.4034790 (Fiedler et al., 2020). Simulated data available from the Dryad Digital Repository upon acceptance of this manuscript.

