## Supplementary for "Restoration ecologists might not get what they want: Global change shifts trade-offs among ecosystem functions"

**S1: Description of the plant module of ModEST** 2

**S2: Model parameterisation and validation** 11

Table S2.1: Plant parameters. 12

Table S2.2: Hydrological and nutrient parameters. 14

Figure S2.1: Calibration results. 15

Figure S2.2: Validation results. 16

**S3: Simulation experiments** 19

S3.1: Processing of weather data from Pingelly 19

Figure S3.1: Plant heights of the one-year old eight species in the Ridgefield experiment. 19

Table S3.1: Median annual and seasonal climate trends. 19

Figure S3.2: Current (left) and future (right) climate diagrams for Pingelly. 20

Figure S3.3: Relative plant cover over time for the 8-species mixture under current climatic conditions. 20

**S4: Supporting results** 22

Figure S4.1: Ecosystem functioning for each planted species richness under current (white boxplots) and different future climatic conditions (boxplots in different shades of grey in the order: RCP 26, 45, 60, and 85). 22

Figure S4.2: Realised richness for each planted richness for current (white) and future conditions (grey). 23

Figure S4.3: Variability of ecosystem functioning under current conditions. 24

Figure S4.4: CWM trait values across all planted species scenarios for current (white) and future conditions (grey). 25

Figure S4.5: CWM trait values for each planted richness for current (white) and future conditions (grey). 26

Figure S4.6: Comparison of different multifunctionality indices across the different species richness and climate change scenarios (white: current, grey: future). 27

### S1: Description of the plant module of ModEST

As described in the main text, the plant module is mainly based on LPJ and LPJmL (Schaphoff et al., 2017; Sitch et al., 2003; Smith et al., 2014) and local processes as described for an individual-based plant model by May et al. (2009). The plant module simulates the life cycle of individual woody plants placed in the landscape, their dynamic below- and aboveground carbon and nitrogen pools as well as structural components based on plant traits (see Table S2.1) and abiotic conditions. As this module is a combination of several literature sources, a few adaptations or corrections to the processes described therein and a new mortality process, we decided to fully describe the plant module in the following.

This module is fully coupled to a spatially explicit hydrological and nutrient module (see description of both modules in the main text), meaning that soil water and nutrient fluxes are influenced by individual plants, and vice versa. If not stated otherwise, all plant processes are calculated on a daily basis for each individual plant in a random order to avoid advantages to first individuals if order was maintained. This module has closed boundary conditions affecting dispersal, resource acquisition and litter accumulation. Interaction with soil resources in grid cells are proportional to the individual's intersected crown area which we assume to be their zone of influence. The processes are executed in the order we describe them.

#### *Phenology*

In this study, woody plants were evergreen. In this module, however, woody plants could potentially be distinguished between evergreen, winter deciduous, and drought deciduous phenology types as described in Sitch et al. (2003). Phenology *phen* (ranging between 0 and 1) varies between fully shed and fully covered with leaves and affects photosynthesis, transpiration and respiration. Evergreen plants retain a constant full leaf coverage over the year. Phenology of drought deciduous depends on a threshold of water supply and water demand (see transpiration for $w_{lim}$), whereas phenology of winter deciduous is a daily updated fraction of current growing degree days based on a species-specific temperature level (GDD) and the maximum GDD for full leaf cover ($full_{GDD}$).

In the case of senescence, 50 % of the optimal leaf nitrogen content will be allocated to the plant's nitrogen storage but only up to its maximum nitrogen storage capacity (Smith et al., 2014). The shed leaves will be added to the aboveground litter pool (proportionally to covered grid cells).

| | *Phenology status* | *Senescence* |
|---|---|---|
| *Evergreen* | $phen = 1.0$ | - |
| *Winter deciduous* | $phen_t = min\left\{1.0, \frac{GDD_t}{full_{GDD}}\right\}$ | The first time if temperature drops below 5.0 °C in autumn. |
| *Drought deciduous* | $hen = \begin{cases} 1.0, & w_{lim} \geq 0.35 \\ 0.0, & w_{lim} < 0.35 \end{cases}$ | $w_{lim} < 0.35$ |

#### *Photosynthesis*

Photosynthesis is based on Sitch et al. (2003), Smith et al. (2014) and Schaphoff et al. (2018) and calculates gross primary production $GPP$ [kgC · day$^{-1}$] based on light-limited $J_E$ [kgC · hour$^{-1}$] and Rubisco-limited photosynthesis $J_C$ [kgC · hour$^{-1}$]:

$$GPP = \frac{J_E + J_C - \sqrt{(J_E + J_C)^2 - 4 \cdot \theta \cdot J_E \cdot J_C}}{2 \cdot \theta} \cdot daylength \quad (1)$$

with $daylength$ as daylight hours [hours] and $\theta = 0.7$, a shape parameter that describes the co-limitation of light and Rubisco activity [dimensionless].

Daylength is simulated as:

$$daylength = \frac{24}{\pi} \cdot \arccos\left(-\frac{\sin\left(\varphi \cdot \frac{\pi}{180}\right) \cdot \sin\delta}{\cos\left(\varphi \cdot \frac{\pi}{180}\right) \cdot \cos\delta}\right) \tag{2}$$

with $\varphi$ giving the latitude of the site [rad] and $\delta$ the solar declination [rad]:

$$\delta = 0.4093 \cdot \sin(\frac{2\pi}{365} \cdot day - 1.405) \tag{3}$$

with $day$ as day number of the year.

$J_E$ depends on absorbed photosynthetically active radiation $APAR$, daylength, and air temperature.

$$J_E = c_1 \cdot \frac{APAR}{daylength} \tag{4}$$

$$c_1 = \alpha_{C3} \cdot T_{Stress} \cdot \left(\frac{p_i - \Gamma_*}{p_i + 2\Gamma_*}\right) \tag{5}$$

with $\alpha_{C3} = 0.08$ as the intrinsic quantum efficiency for $CO_2$ uptake in $C_3$ plants [dimensionless], $T_{Stress}$ as a species-specific temperature inhibition function [dimensionless] (Schaphoff et al., 2018), $p_i$ as leaf internal partial pressure of $CO_2$ [Pa], and $\Gamma_*$ as the photorespiratory $CO_2$ compensation point [Pa].

$$\begin{aligned} T_{Stress} &= \frac{1.0}{1 + e^{k_1 \cdot (k_2 - T_{air})}} \cdot \left(1.0 - 0.01 \cdot e^{k_3 \cdot (T_{air} - T_3)}\right) \\ k_1 &= 2.0 \cdot \frac{\ln\left(\frac{1.0}{0.99} - 1.0\right)}{T_1 - T_2} \\ k_2 &= 0.5 \cdot (T_1 + T_2) \\ k_3 &= \frac{\ln\frac{0.99}{0.01}}{T_4 - T_3} \end{aligned} \tag{6}$$

with $T_1$ to $T_4$ as species-specific temperature limits [°C], $T_{air}$ as air temperature [°C].

$$p_i = \lambda \cdot p_a \tag{7}$$

with $p_a$ as ambient pressure of $CO_2$ [Pa], $\lambda$ as parameter describing the ratio of the intercellular to the ambient $CO_2$ concentration [dimensionless].

$$\Gamma_* = \frac{[O_2]}{2\tau} \tag{8}$$

with $[O_2] = 20900$ as partial pressure of $O_2$ [Pa], $\tau$ as specificity factor that reflects the ability of Rubisco to discriminate between $CO_2$ and $O_2$ [dimensionless].

$$\tau = \tau_{25} \cdot Q_{10_\tau}^{(T_{air} - 25) \cdot 0.1} \tag{9}$$

with $\tau_{25} = 2600$ as τ at 25 °C [dimensionless], $Q_{10_\tau} = 0.57$ as temperature sensitivity factor [dimensionless], $T_{air}$ as mean daily air temperature [°C].

$APAR$ $[\frac{\text{kgC}}{\text{day}}]$ is the fraction of incoming net photosynthetically active radiation PAR $[\frac{\text{kgC}}{\text{m}^2\text{day}}]$ that is absorbed by the unshaded plant fractional coverage $FPC_{unshaded}$ [$\text{m}^2$] given the daily phenological status [dimensionless].

$$PAR = PAR \cdot FPC_{unshaded} \cdot phen$$
$$PAR = 0.5 \cdot c_q \cdot c_{mass} \cdot R_a \qquad (10)$$

with $c_q = 4.6 \times 10^{-6}$ $[\frac{\text{J}}{\text{mol}}]$ conversion factor from J to mol for solar radiation at 550 nm, $c_{mass} = 12.0107 \cdot 10^{-3}$ $[\frac{\text{kg}}{\text{mol}}]$ as atomic mass of carbon, and $R_a$ as extraterrestrial radiation $[\frac{\text{J}}{m^2 \cdot \text{day}}]$.

$FPC_{unshaded}$ of an individual is calculated from the leaf area index $LAI$ [dimensionless] and the individual unshaded crown area $CA_{unshaded}$ [$\text{m}^2$] which is the crown area that is not overlapped by the crowns of larger surrounding individuals.

$$FPC_{unshaded} = (1 - e^{-0.5 \cdot LAI}) \cdot CA_{unshaded}$$
$$LAI = \frac{C_{leaf} \cdot SLA}{CA} \qquad (11)$$

with $C_{leaf}$ as dynamic carbon mass in the leaves [kg], and $SLA$ as species-specific leaf area $[\frac{m^2}{kg}]$.

$J_C$ is calculated as a function of maximum Rubisco capacity $V_m$ $[\frac{\text{kgC}}{\text{day}}]$.

$$J_C = \frac{c_2 \cdot V_m}{24} \qquad (12)$$

$$c_2 = \frac{p_i - \Gamma_*}{p_i + K_c \cdot \left(1.0 + \frac{[O_2]}{K_o}\right)} \qquad (13)$$

with $K_c = 30$ and $K_o = 30000$ as Michaelis-Menten constants at 25 °C [Pa] and other symbols as defined previously.

$$V_m = \frac{1}{b} \cdot \frac{c_1}{c_2} \cdot [(2 \cdot \theta - 1) \cdot s - (2 \cdot \theta \cdot s - c_2) \cdot \sigma] \cdot APAR \cdot n_{lim} \qquad (14)$$

with $b = 0.015$ as leaf respiration as fraction of $V_m$ for $C_3$ plants, $n_{lim}$ as nitrogen limitation factor (see description of *Nitrogen uptake*) which is a simplification of nitrogen limited Rubisco activity as described in Smith et al. (2014).

$$s = \frac{24}{daylength} \cdot b \qquad (15)$$

$$\sigma = \sqrt{1.0 - \frac{c_2 - s}{c_2 - \theta \cdot s}} \qquad (16)$$

Stomatal conductance $g_c$ [$\text{mm} \cdot \text{s}^{-1}$] is calculated as:

$$g_c = \frac{1.6 \cdot NPP_{mm/s}}{c_a \cdot (1.0 - \lambda)} + g_{min} \qquad (17)$$

with as net daytime primary production $NPP_{mm/s}$ [$\text{mm} \cdot \text{s}^{-1}$], $c_a$ as ambient mole fraction of $CO_2$, and $g_{min}$ as species-specific minimum canopy conductance [$\text{mm} \cdot \text{s}^{-1}$] that occurs due to processes other than photosynthesis.

$$c_a = \frac{p_a}{p} \tag{18}$$

with $p = 1 \cdot 10^5$ [Pa] as atmospheric pressure, and $p_a$ as defined previously.

$NPP_{mm/s}$ is calculated using the ideal gas equation:

$$NPP_{mm/s} = \frac{NPP}{conv_{DtoS}} \cdot \frac{1}{c_{mass}} \cdot \bar{R} \cdot \frac{T_{air} + 273.15}{p} \cdot 1000 \tag{19}$$

with $conv_{DtoS} = 3600\ \mathrm{s} \cdot \mathrm{hour}^{-1} \cdot daylength$ as conversion factor from daylight hours to seconds, $\bar{R} = 8.314$ [$\mathrm{m}^3 \cdot \mathrm{Pa} \cdot \mathrm{K}^{-1} \cdot \mathrm{mol}^{-1}$] as ideal gas constant.

Net primary productivity $NPP$ [kgC · day$^{-1}$] results from $GPP$ (see eqn. 1) after leaf respiration $R_{leaf}$ [kgC · day$^{-1}$]:

$$NPP = GPP - R_{leaf} \tag{20}$$

$$R_{leaf} = \frac{daylength}{24} \cdot b \cdot V_m \tag{21}$$

As described in Sitch et al. (2003), under non-water-stressed conditions $\lambda = 0.8$ for $C_3$ plants, and therefore the resulting $g_c$ is the potential stomatal conductance which will be used in the *Transpiration* process to assess the potential water demand. If the water demand cannot be fulfilled by the water supply $W_{sup}$ (see *Transpiration*), equations involving water demand $W_{dem}$ (see *Transpiration*), $g_c$, and $NPP$ will to be solved again but this time simultaneously to obtain values for $g_c$ and $\lambda$ that meet $W_{sup}$. This ultimately leads into a down-regulation of the photosynthesis and thus a decreased $GPP$.

***Nitrogen uptake***

Nitrogen uptake is simplified after the description of Smith et al. (2014). An individual's potential nitrogen demand $N_{pot,dem}$ [kgN] is the sum of the nitrogen demand for the leaves $N_{leaf,dem}$, sapwood $N_{sap,dem}$ and roots $N_{root,dem}$ as well as demand for the storage of labile nitrogen $N_{store,dem}$.

$$N_{pot,dem} = N_{leaf,dem} + N_{sap,dem} + N_{root,dem} + N_{store,dem} \tag{22}$$

Demand for each tissue $N_{tissue,dem}$ is based on the optimal carbon to nitrogen ratio $CN_{tissue}$and the current carbon mass of the tissue $C_{tissue}$ [kgC].

$$N_{tissue,dem} = \frac{C_{tissue}}{CN_{tissue}} \tag{23}$$

Demand for nitrogen storage pool $N_{store,dem}$ is the difference between current storage $N_{store}$ and the maximum nitrogen storage capacity $N_{store,max}$.

$$N_{store,dem} = N_{store,max} - N_{store} \tag{24}$$

$N_{store,max}$ is related to the current sapwood carbon $C_{sap}$, leaf nitrogen $N_{leaf}$ and carbon $C_{leaf}$, as well as $k_{store}$ a constant which is set to 0.05 for evergreen woody and 0.15 for deciduous woody plants.

$$N_{store,max} = k_{store} \cdot C_{sap} \cdot \frac{N_{leaf}}{C_{leaf}} \tag{25}$$

Actual nitrogen demand $N_{act,dem}$ is limited by maximal possible N uptake $N_{max}$. If the potential demand cannot be fulfilled by $N_{max}$, tissue demand will be adjusted accordingly.

$$N_{act,dem} = \begin{cases} N_{pot,dem}, & N_{pot,dem} < N_{max} \\ N_{max}, & N_{pot,dem} \geq N_{max} \end{cases} \tag{26}$$

$N_{max}$ depends on the plant available soil nitrogen $f_{avN}$, soil temperature $f_{T,soil}$, current N:C status of the plant $f_{NC}$, and root carbon mass $C_{root}$.

$$N_{max} = 2.0 \cdot up_{N,root} \cdot f_{avN} \cdot f_{T,soil} \cdot f_{NC} \cdot C_{root} \tag{27}$$

with $up_{N,root} = 2.8 \cdot 10^{-3} \frac{\mathrm{kgN}}{\mathrm{kgC\ day}}$ for woody plant species.

$$f_{avN} = 0.05 + \frac{N_{soil}}{N_{soil} + k_{max} \cdot fc \cdot z_{soil} \cdot CA} \tag{28}$$

with $N_{soil}$ as the nitrate $NO3$ [kg] and ammonium $NH4$ [kg] content in the zone of influence of the individual plant (output from nutrient module); $k_{max} = 1.48 \cdot 10^{-3} \frac{\mathrm{kgN}}{\mathrm{m^3}}$ as the half-saturation concentration for N uptake for woody plant species; $fc$ field capacity of the soil $[\frac{\mathrm{m^3}}{\mathrm{m^3}}]$, $z_{soil}$ soil depth in which the plant is rooting [m]; and crown area $CA$ [$\mathrm{m^2}$] .

$$N_{soil} = \sum_{C}^{nC's} \sum_{L}^{nL's} (NO3_L + NH4_L) \cdot root_L \cdot relCA_C \tag{29}$$

with $relCA_C$ being the relative plant's crown area intersected for each grid cell $C$, and $root_L$ being the relative fraction of root mass per soil layer $L$.

$$f_{T,soil} = 0.0326 + 0.00351 \cdot {T_{av,soil}}^{1.652} - \left(\frac{T_{av,soil}}{41.748}\right)^{7.19} \tag{30}$$

As soil temperature $T_{soil}$ [°C] (from nutrient module, see Kemanian et al., 2011) is different across soil layers $L$ and grid cells $C$, an average soil temperature $T_{av,soil}$ per plant individual is calculated based on the fraction of roots per soil layer $root_L$ [–] and the relative intersected crown area per grid cell $relCA_C$ [–].

$$T_{av,soil} = \sum_{C}^{nC's} \sum_{L}^{nL's} T_{soil,L} \cdot root_L \cdot relCA_C \tag{31}$$

$$f_{NC} = \frac{x_1}{x_2}$$

$$x_1 = \frac{N_{leaf} + N_{root}}{C_{leaf} + C_{root}} - \frac{1}{CN_{leaf,min}} \tag{32}$$

$$x_2 = \frac{2.0}{CN_{leaf,max} + CN_{leaf,min}} - \frac{1.0}{CN_{leaf,min}}$$

with $CN_{leaf,max}$ and $CN_{leaf,min}$ as the maximum and minimum bounds for leaf C:N, respectively.

Nitrogen is taken up according to the actual N demand of root, sapwood and leaf tissue and the storage pool. If there is only enough available N to meet the demand of the tissue, demand for the storage will not be fulfilled. If there is not enough to meet tissue N demand, N from the storage pool will be used. If there still is not enough available N to meet the tissue's demand, photosynthesis will be re-calculated with limiting term $n_{lim}$ (see *Photosynthesis*).

$$n_{lim} = min\left\{ 1.0, \frac{N_{av,soil} + N_{store}}{N_{leaf,dem} + N_{sap,dem} + N_{root,dem}} \right\} \tag{33}$$

***Transpiration***

Transpiration $E_T$ is based on the approach by Sitch et al. (2003) and Schaphoff et al. (2018) and is assumed to be the minimum between water demand $W_{dem}$ and water supply $W_{sup}$.

$$E_T = min\{W_{dem}, W_{sup}\} \tag{34}$$

If $W_{sup}$ cannot meet $W_{dem}$, water limitation $w_{lim}$ will be calculated and considered in the allocation (see *Allocation*), as well as stomatal conductance $g_c$ will be re-calculated given $W_{sup}$ (see *Photosynthesis*). The amount of water transpired is calculated proportional to the covered grid cells and the root fraction in the different soil layers.

$$w_{lim} = min\left\{1.0, \frac{W_{sup}}{W_{dem}}\right\} \tag{35}$$

$W_{dem}$ [mm] is calculated based on potential evapotranspiration $E_{pot}$ (from hydrological module, see Tietjen et al., 2009) and fractional plant cover $FPC$ (see $FPC_{unshaded}$ in *Photosynthesis* but with full crown area $CA$) and stomatal conductance $g_c$ (see *Photosynthesis*).

$$W_{dem} = E_{pot} \cdot FPC \cdot \frac{\alpha_m}{1 + \frac{g_m}{g_c}} \tag{36}$$

with maximum Priestley-Taylor coefficient $\alpha_m = 1.1$ (Monteith, 1995) and conductance scaling factor $g_m = 5.0$ (Sitch et al., 2003).

$W_{sup}$ [mm] is calculated based on a species-specific maximum water transport capacity $e_{max}$ [$\frac{mm}{\text{day}}$], fractional plant cover $FPC$, relative water content available for the plant $W_{av}$, phenology $phen$, and a scaling term based on the current root to leaf mass ratio allowing more or less water supply with deviating species-specific leaf mass to root mass ratio $LM/RM$.

$$W_{sup} = e_{max} \cdot FPC \cdot W_{av} \cdot phen \cdot \frac{C_{root} \cdot LM/RM}{C_{leaf}} \tag{37}$$

Relative water content available for the plant $W_{av}$ is the difference between the current relative water content $W$ (output from hydrological module) and water content at which plant starts to wilt $WP$ as a fraction of field capacity $fc$ of the soil.

$$W_{av} = \sum_{C}^{nC's} \sum_{L}^{nL's} \left(\frac{W_L - WP}{fc}\right) \cdot root_L \cdot relCA_C \tag{38}$$

with $relCA_C$ being the relative plant's crown area intersected for each grid cell $C$, and $root_L$ being the relative fraction of root mass per soil layer $L$.

***Respiration***

Respiration is modelled as described in Sitch et al. (2003) and updates described in Schaphoff et al. (2018). An individual's total respiration $R$ [kgC] is a sum of maintenance leaf $R_{leaf}$ (see photosynthesis), sapwood $R_{sap}$ [kgC] and root respiration $R_{root}$ [kgC] as well as growth respiration $R_{growth}$ [kgC].

$$R = R_{leaf} + R_{sap} + R_{root} + R_{growth} \tag{39}$$

Maintenance respiration depends on tissue-specific carbon to nitrogen ratio $CN_{tissue}$, air or soil temperature $T_x$ for sapwood or root respiration, respectively, tissue's biomass $C_{tissue}$ and phenology $phen$ for root respiration.

$$R_{sap} = r_{coef} \cdot k \cdot \frac{C_{sap}}{CN_{sap}} \cdot A_x \tag{40}$$

$$R_{root} = r_{coef} \cdot k \cdot \frac{C_{root}}{CN_{root}} \cdot A_x \cdot phen \tag{41}$$

with $r_{coef}$ as species-specific respiration coefficient, $k = 0.095218$ and Arrhenius temperature-respiration function.

$$A_x = e^{308.56 - \left(\frac{1}{56.02} - \frac{1}{T_x + 46.02}\right)} \tag{42}$$

with x as air or average soil temperature ($T_{av,soil}$, see *Nitrogen Uptake*) for sapwood or roots, respectively.

Growth respiration is a fixed proportion of 25 % of the remaining GPP after subtracting maintenance respiration.

$$R_{growth} = r_{gr} \cdot (GPP - R_{leaf} - R_{sap} - R_{root}) \tag{43}$$

with $r_{gr} = 0.25$.

As a result, net primary production $NPP = GPP - R$ will be calculated.

***Reproduction***

Reproduction is modelled as described in Sitch et al. (2003). 10 % of daily NPP goes into reproductive carbon mass $m_{rep,d}$ and will be summed up over one year $m_{rep,y}$ [kgC]. The remaining net primary productivity $NPP_{rem}$ [kgC · day$^{-1}$] will be calculated as follows:

$$NPP_{rem} = NPP - m_{rep,d} \tag{44}$$

***Allocation***

Allocation is based on Sitch et al. (2003) and updates as described in Smith et al. (2014). Accordingly, GPP after respiration and reproduction, $NPP_{rem}$, will be allocated to above- and below-ground carbon pools, i.e. leaves, fine roots and sapwood by satisfying four allometric relationships (eqns. 47 to 50).

$$NPP_{rem} = \Delta C_{leaf} + \Delta C_{root} + \Delta C_{sap} \tag{45}$$

$$LA = LtS \cdot SA$$
$$SA = \frac{C_{sapwood}}{WD \cdot H} \tag{46}$$

with $LA$ as average individual leaf area [m$^2$], $SA$ as sapwood cross sectional area [m$^2$], $LtS$ as a species-specific ratio between leaf area to sapwood area, $WD$ as species-specific wood density [$\frac{\text{kgC}}{\text{m}^3}$], and $H$ as plant height [m].

$$\Delta C_{leaf} = LtR_{scal} \cdot \Delta C_{root} \tag{47}$$

with $LtR_{scal}$ as leaf mass to root mass constant depending on species-specific $LtR$ as well as water or nutrient limitation leading to greater allocation to the roots driven by the most limiting resource.

$$LtR_{scal} = \begin{cases} LtR, & NPP \leq 0 \\ LtR \cdot min\{w_{lim}, n_{lim}\}, & NPP > 0 \end{cases} \tag{48}$$

$$H = a_2 \cdot D^{a_3}$$
$$D = \left(\frac{4 \cdot (C_{heartwood} + C_{sapwood})}{\pi \cdot WD \cdot a_2}\right)^{\frac{1}{a_3+2}} \tag{49}$$

with $D$ as stem diameter [m] as well as $a_2$ and $a_3$ as species-specific allometric parameters.

$$CA = \begin{cases} a_1 \cdot D^{k_{rp}}, & CA < CA_{max} \\ CA_{max}, & CA \geq a_1 \cdot D^{k_{rp}} \end{cases} \quad (50)$$

with CA as crown area constrained by a species-specific maximum crown area $CA_{max}$ and species-specific allometric parameter $a_1$ as well as $k_{rp} = 1.6$.

If there is negative NPP, this results in negative $\Delta C_{leaf}$, $\Delta C_{root}$ and potentially negative $\Delta C_{\mathrm{sap}}$. In this case, leaf and root carbon will be lost to the aboveground or belowground litter pool, and sapwood carbon will be transformed to heartwood carbon mass. Total nitrogen in the lost tissue $N_{turnover}$, which is calculated from $\Delta C_{leaf}$, $\Delta C_{root}$ and $\Delta C_{\mathrm{sap}}$ and the species- and tissue-specific $CN_{tissue}$, is moved to the plant's labile nitrogen storage which is constrained by the maximum possible nitrogen storage $N_{store,max}$. Excess nitrogen will be equally added to the above- and belowground nitrogen litter pools.

$$N_{turnover} = \sum_{tissue}^{n} \frac{|\Delta C_{tissue}|}{CN_{tissue}}, \Delta C_{tissue} < 0 \quad (51)$$

$$N_{store_t} = min\{N_{store_{t-1}} + N_{turnover}, N_{store,max}\} \quad (52)$$

***Dispersal & Establishment***

The dispersal routine is based on May et al. (2009) and is executed once a year on July 1st or for leap years on July 2nd (for southern hemisphere). Number of seeds will be determined by the individual's reproductive carbon mass allocated over the year $m_{rep,y}$ divided by the species-specific mass of one seed *seedMass* [kg]. Distance [m] is drawn from a log normal distribution with species-specific mean *meanDisp* and standard deviation *sdDisp* of dispersal distance which resembles the species-specific dispersal kernel. Dispersal direction [deg] for each seed is drawn from a uniform distribution.

The saplings establish as an individual plant at the determined location with a height of 10 cm and a species-specific initial leaf area index $LAI_{sap}$ (to obtain initial leaf carbon, eqn. 12) (i) with a species-specific germination probability $p_{germ}$, (ii) if the target position is not covered by another individual assuming too limited light resources (after May et al., 2009), and (iii) with a species-specific establishment probability $p_{estab}$ which equals $m_{seed}$ taking into account increased probability of establishment with higher endosperm resources (Hallett et al., 2011). Seeds that do not establish will be transferred to the aboveground carbon litter pool.

***Mortality***

Mortality events due to resource limitations take place the last day of the year for individuals older than one year. As plant growth depends on resource availability, individual plants die if the ratio between current and previous year's total plant biomass exceeds a species-specific mortality threshold *tmort*. Except for the heartwood pool, which remains as standing biomass, all other individual's above- and belowground carbon and nitrogen pools will be transferred to the respective litter pools of those grid cells and soil layers the individual's crown area intersect with – proportional to the covered crown area per grid cell and the fraction of roots per soil layer $froot_{Lx}$.

*References*

Hallett, L.M., Standish, R.J., Hobbs, R.J., 2011. Seed mass and summer drought survival in a Mediterranean-climate ecosystem. Plant Ecol. 212, 1479–1489. https://doi.org/10.1007/s11258-011-9922-2

Kemanian, A.R., Julich, S., Manoranjan, V.S., Arnold, J.R., 2011. Integrating soil carbon cycling with that of nitrogen and phosphorus in the watershed model SWAT: Theory and model testing. Ecol. Modell. 222, 1913–1921. https://doi.org/10.1016/j.ecolmodel.2011.03.017

May, F., Grimm, V., Jeltsch, F., 2009. Reversed effects of grazing on plant diversity: The role of below-ground competition and size symmetry. Oikos 118, 1830–1843. https://doi.org/10.1111/j.1600-0706.2009.17724.x

Monteith, J.L., 1995. Accomodation Between Transpiring Vegetation and the Convective Boundary-Layer. J. Hydrol. 166, 251–263. https://doi.org/10.1016/0022-1694(94)05086-d

Schaphoff, S., von Bloh, W., Rammig, A., Thonicke, K., Biemans, H., Forkel, M., Gerten, D., Heinke, J., Jägermeyr, J., Knauer, J., Langerwisch, F., Lucht, W., Müller, C., Rolinski, S., Waha, K., 2017. LPJmL4 - a dynamic global vegetation model with managed land: Part I - Model description. Geosci. Model Dev. Discuss. 1–59. https://doi.org/10.5194/gmd-2017-145

Sitch, S., Smith, B., Prentice, I.C., Arneth, a., Bondeau, a., Cramer, W., Kaplan, J.O., Levis, S., Lucht, W., Sykes, M.T., Thonicke, K., Venevsky, S., 2003. Evaluation of ecosystem dynamics, plant geography and terrestrial carbon cycling in the LPJ dynamic global vegetation model. Glob. Chang. Biol. 9, 161–185. https://doi.org/10.1046/j.1365-2486.2003.00569.x

Smith, B., Wärlind, D., Arneth, A., Hickler, T., Leadley, P., Siltberg, J., Zaehle, S., 2014. Implications of incorporating N cycling and N limitations on primary production in an individual-based dynamic vegetation model. Biogeosciences 11, 2027–2054. https://doi.org/10.5194/bg-11-2027-2014

Tietjen, B., Zehe, E., Jeltsch, F., 2009. Simulating plant water availability in dry lands under climate change: A generic model of two soil layers. Water Resour. Res. 45, 1–14. https://doi.org/10.1029/2007WR006589

### S2: Model parameterisation and validation

Whenever possible, the model parameters of ModEST were parameterized according to values from measurements in Ridgefield, literature, or databases (see Tables S2.1 and S2.2). Parameters with unknown values were calibrated. For *E. astringens* additional known parameters through measurements (i.e. LM/RM, $a_2$, $a_3$) were included in the calibration process, as they proved important for a successful calibration.

For the actual calibration, ModEST was initialised for conditions found in Ridgefield in 2011. We used the same plot size, the coordinates of individual plants, and their measured heights. Soil conditions of these plots were mimicked in terms of the measured nitrate and ammonium content in the soil, bulk density, and clay content (Perring et al., 2012). We run simulations from 2011 to 2016 using weather time series dataset from Ridgefield where available (2013 – 2016) and filled the gap in the first years with the nearby weather station in Pingelly (2011 – 2012; Bureau of Meteorology, 2019). The modelled landscape was flat with 50 and 150 cm in depth for the first and the second soil layer, respectively. Since monoculture data was only available for *E. loxophleba*, we calibrated the parameters of each plant species consecutively. We first used a monoculture plot for *E. loxophleba* (for which soil parameters were calibrated at the same time), followed by a two-species mixture plot for *E. astringens*, and then four-species mixtures plots for *A. acuminata, A. microbotrya, B. sessilis, H. lissocarpha, C. quadrifidus, C. phoeniceus* (see Perring et al., 2012 for information on the mixtures). For each plant species, we compared yearly simulated mean aboveground alive biomass, mean plant height and individual counts with measured data from the plots. We compared estimates for August, since this is when these variables were measured in the experiment, on an annual basis from 2011 until 2014 (for individual counts until 2016). Soil moisture time series data were only available for the *E. loxophleba* plot, therefore we calibrated soil parameters for this plot using daily time series data of the first soil layer (Fig. S2.1). Calibrated parameters were obtained by a Bayesian approach implemented in the R package BayesianTools (Hartig et al., 2019) using 100,000 iterations, 3 chains, and the DEzs sampler. Input parameter ranges (priors) can be found in Tables S2.1 and S2.2. Optimisation across the abovementioned four output variables was measured by a root square sum error between measured $x_m$ and simulated data $x_s$ normalized by the standard deviation $\sigma_m$ of the measured data at each time of the measurement $i$ (nRSSE). For each parameter, we used the maximum probable value of its calibrated posterior distribution.

For model validation (see also in the main text *Model parameterization and validation*, Fig. S2.2), we used the same settings as described for model calibration. Soil water and nutrient dynamics could not be validated due to a lack of empirical data.

**Table S2.1: Plant parameters.** For meaning of the parameters see plant module description (Supplementary S1). Highlighted parameters in grey were focal traits in this study (Table 2 in main text). EL: *Eucalyptus loxophleba* ssp. *loxophleba,* EA: *E. astringens,* AA: *Acacia acuminata,* AM: *A. microbotrya,* BS: *Banksia sessilis,* HL: *Hakea lissocarpha,* CQ: *Calothamnus quadrifidus,* CP: *Callistemon phoeniceus.*

| Parameter | Unit | Reference to Supp. S1 | Value for each woody evergreen plant species | | | | | | | | Source |
|---|---|---|---|---|---|---|---|---|---|---|---|
| | | | EL | EA | AA | AM | CP | CQ | BS | HL | |
| $T_1$ | °C | eqn. 6 | -4 | | | | | | | | Sitch et al., (2000), temperate needle/broad-leaved evergreen species |
| $T_2$ | °C | | 20 | | | | | | | | |
| $T_3$ | °C | | 30 | | | | | | | | |
| $T_4$ | °C | | 42 | | | | | | | | |
| SLA | $m^2\ kg^{-1}$ | eqn. 12 | 4.14 | 3.3 | 6.4 | 6.17 | 3.18 | 3.35 | 5.42 | 4.07 | Fiedler et al., (unpublished) |
| $g_{min}$ | $mm\ s^{-1}$ | eqn. 18 | 0.64 | 0.991 | 0.992 | 0.999 | 0.301 | 0.957 | 0.302 | 0.326 | calibrated between 0.3 and 1.0 (default = 0.5) based on Schaphoff et al., (2017) |
| $CN_{leaf}$ | - | eqn. 24 | 32.4 | 53.4 | 16.9 | 20.4 | 48.7 | 36.7 | 53 | 72.4 | Fiedler et al., (unpublished) |
| $CN_{sap}$ | - | eqns. 24, 41 | 100 | 172.6 | 41.9 | 50.8 | 81.9 | 86.3 | 112 | 92.9 | Fiedler et al., (unpublished) |
| $CN_{root}$ | - | eqns. 24, 42 | 84.6 | 123.18 | 34.6 | 44.6 | 100.6 | 49.9 | 73.6 | 54.75 | Perring et al., (unpublished) |
| $k_{store}$ | - | eqn. 26 | 0.05 | | | | | | | | Smith et al., (2014) |
| $root_{L1}$ | - | eqns. 30, 32 (also used in hydrological module) | 0.023[*1] | 0.012[*1] | 0.48[*1] | 0.027[*1] | 0.999[*2] | 0.201[*2] | 0.387[*2] | 0.011[*2] | *1: calibrated between 0 and 0.5 (default = 0.5) for trees<br>*2: calibrated between 0 and 1.0 (default = 0.5) for shrubs |
| $CN_{leaf,min}$ | - | eqn. 33 | 29.1 | 46.8 | 13.9 | 19.8 | 40.5 | 34.5 | 38.3 | 66.2 | Fiedler et al., (unpublished) |
| $CN_{leaf,max}$ | - | eqn. 33 | 34.7 | 58.6 | 20.9 | 21 | 61.4 | 38.9 | 71.1 | 83.7 | Fiedler et al., (unpublished) |
| $e_{max}$ | $mm\ day^{-1}$ | eqn. 38 | 12.953 | 7.527 | 5.092 | 5.544 | 5.02 | 5.17 | 13.328 | 12.169 | calibrated between 5 and 15 (default = 5) based on Sitch et al., (2003) |
| LM/RM | - | eqn. 38 | 0.92[*1] | 3.933[*2] | 0.43[*1] | 0.47[*1] | 0.96[*1] | 1.73[*1] | 0.8[*1] | 1.54[*1] | *1: Perring et al., (unpublished)<br>*2: calibrated between 0.9 and 4.26 (default = 2.06) based on Perring et al., (unpublished) |
| WP | - | eqn. 39 | 0.102 | 0.148 | 0.085 | 0.148 | 0.066 | 0.1 | 0.143 | 0.093 | calibrated between 0.05 and 0.15 (default = 0.1) based on Rawls et al., (1992) for loamy sand |
| $r_{coef}$ | - | eqns. 41, 42 | 0.057 | 0.015 | 0.069 | 0.05 | 0.759 | 0.467 | 0.407 | 1.192 | Calibrated between 0.01 and 1.2 (default = 0,2) based on Schaphoff et al., (2017) |
| LtS | - | eqn. 47 | 8000 | | | | | | | | Sitch et al., (2003) |
| WD | $kgC\ m^{-3}$ | eqns. 47, 50 | 973.791[*1] | 824.946[*1] | 951.424[*1] | 815.68[*1] | 846.363[*1] | 971.899[*2] | 771.1[*1] | 832.5135[*1] | *1: Kattge et al., (2011) |

| Parameter | Unit | Reference to Supp. S1 | Value for each woody evergreen plant species | | | | | | | | Source |
|---|---|---|---|---|---|---|---|---|---|---|---|
| | | | EL | EA | AA | AM | CP | CQ | BS | HL | |
| | | | | | | | | | | | *2: calibrated between 600 and 1000 (default = 800) |
| $a_2$ | - | eqn. 50 | 34.9[*1] | 40.48[*2] | 43.075[*1] | 33.235[*1] | 63.614[*2] | 30.328[*2] | 141.98[*1] | 61.959[*2] | *1: Perring et al., (unpublished)<br>*2: calibrated:<br>EA between 21.7 and 56.6 (default = 39.15) based on Perring et al., (unpublished)<br>CP between 1 and 100 (default = 100), CQ between 10 and 200 (default = 100), HL between 1 and 100 (default = 10) based on pre-tests |
| $a_3$ | - | eqn. 50 | 0.79[*1] | 0.97[*2] | 0.825[*1] | 0.825[*1] | 1.002[*2] | 1.113[*2] | 1.33[*1] | 1.171[*2] | *1: Perring et al., (unpublished)<br>*2: calibrated:<br>EA between 0.56 and 1.08 (default = 0.82) based on Perring et al., (unpublished)<br>CP between 1 and 2 (default = 1.5), CQ between 0.5 and 2 (default = 1), HL between 0.8 and 1.2 (default = 1) based on pre-tests |
| $a_1$ | - | eqn. 51 | 585.485 | 565.529 | 589.553 | 596.315 | 597.588 | 145.158 | 592.695 | 210.474 | calibrated between 50 and 600 (default = 500) based on Schaphoff et al., (2017) |
| maxCA | $m^2$ | eqn. 51 | 100 | 100 | 50 | 50 | 15 | 15 | 40 | 10 | estimated from maximum height (see Perring et al., 2012):<br>maxCA ≈ 6.7 * maximum height |
| seedMass | mg | Dispersal & Establishment | 0.87 | 7.95 | 15.64 | 28.67 | 0.0462 | 0.91 | 36.67 | 23.9 | Kattge et al., 2011 |
| meanDisp | m | Dispersal & Establishment | 13.815 | 13.815 | 1.708 | 1.708 | 1.604 | 4.01 | 4.812 | 1.203 | Relationship between plant height (Perring et al., 2012) and dispersal distance (Thomson et al., 2011) and dispersal mode (Harris & Standish, 2008) |
| sdDisp | m | Dispersal & Establishment | 2 | | | | | | | | assumed |
| $p_{germ}$ | - | Dispersal & Establishment | 0.97 | 0.92 | 1 | 1 | 1 | 0.9 | 0.88 | 0.96 | Kattge et al., (2011) |
| $LAI_{ini}$ | - | Dispersal & Establishment | 5.042 | 5.786 | 1.072 | 5.997 | 1.675 | 2.932 | 1.755 | 2.621 | calibrated between 1 and 6 (default = 3) |
| tmort | - | Mortality | 1.021[*1] | 1.255[*1] | 1[*2] | 1.186[*1] | 1[*2] | 1.134[*1] | 1.446[*1] | 1.542[*1] | *1: calibrated between 0.5 and 2.0 (default = 0.9)<br>*2: assumed |

**Table S2.2: Hydrological and nutrient parameters.** Parameter shown for the soil texture loamy sandy. Parameters used in plant module (PM, Supplementary S1), hydrological module (HM) by Tietjen et al. (2009), and nutrient module (NM) by Kemanian et al. (2011).

| Parameter | Abb. | Reference to submodules | Unit | Value | Source |
|---|---|---|---|---|---|
| Effective suction | $S_f$ | HM: eqn. 4 | mm | 61.3 | Rawls et al., (1992) |
| Saturated hydraulic conductivity | $K_s$ | HM: eqns. 4, 9 | mm h$^{-1}$ | 62.2 | Rawls et al., (1998) |
| Field capacity | fc | HM<br>PM: eqn. 29 | m$^3$ m$^{-3}$ | 0.332 | calibrated between 0.019 and 0.432 (default = 0.3) based on Rawls et al., (1992) |
| Evaporation factor | $a_{ET}$ | HM: eqn. 13 | - | 0.203 | calibrated between 0.01 and 1.0 (default = 1.0) |
| Residual Water Content | rw | HM: eqn. 13 | m$^3$ m$^{-3}$ | 0.08 | calibrated between 0.003 and 0.12 (default = 0.1) |
| Infiltration rate into layer 2 | $F_{L2,frac}$ | HM: eqn. 2 | - | 0.242 | calibrated between 0.01 and 1.0 (default = 0.1) |
| Bare soil infiltration rate into layer 2 | $F_{L2,bare}$ | HM: eqn. 3 | - | 0.8 | estimated depending on K |
| Maximum total infiltration into layer 2 | $F_{L2,max}$ | HM | mm h$^{-1}$ | 1.5 | Rawls et al., (1992) |
| Constant for diffusion coefficient | d | HM: eqn. 9 | - | 0.253 | calibrated between 0.01 and 1.0 (default = 0.01) |
| Bulk density | BD | NM: eqn. 4 | g cm$^{-3}$ | variable | For calibration and validation depending on measured value of the Ridgefield plot (Perring et al. 2012) |
| | | | | 1.26575 0857 | For scenarios based on mean across all treatments with soil texture loamy sand (Perring et al. 2012) |
| Clay content | $c_{clay}$ | NM: eqns. 3c, 4, 8 | m$^3$ m$^{-3}$ | variable | For calibration and validation depending on measured value of the Ridgefield plot (Perring et al. 2012) |
| | | | | 0.06 | For scenarios based on mean across all treatments with soil texture loamy sand (Perring et al. 2012) |
| Daily nitrogen deposition | $n_{dep}$ | NM | g ha$^{-1}$ year$^{-1}$ | 4.93 | Dentener et al., (2006) |

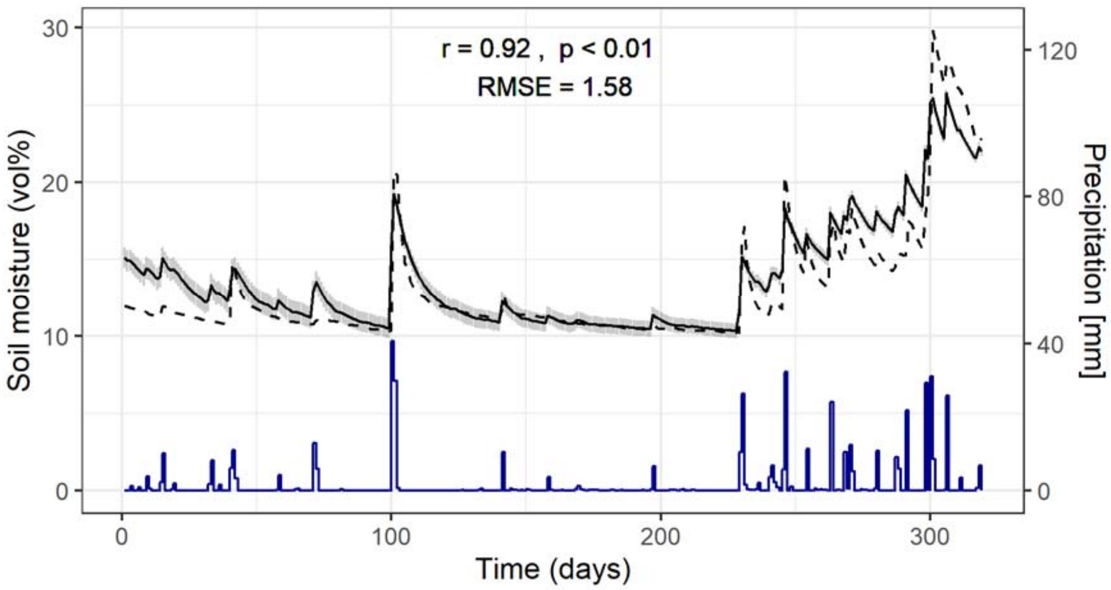


**Figure S2.1: Calibration results.** Shown are measured (dashed line), mean simulated moisture of the first soil layer (solid line) and forcing rainfall (blue line). Measured dynamics show measurement for an annual cycle of a *E. loxophleba* monoculture with loamy sand. Standard deviation shown for simulated results represent 10 model repetitions accounting for random plant height initialisation of neighboring individuals. Shown are Spearman's rank correlation coefficient *r*, corresponding p-value as well as the root mean square error *RMSE*.

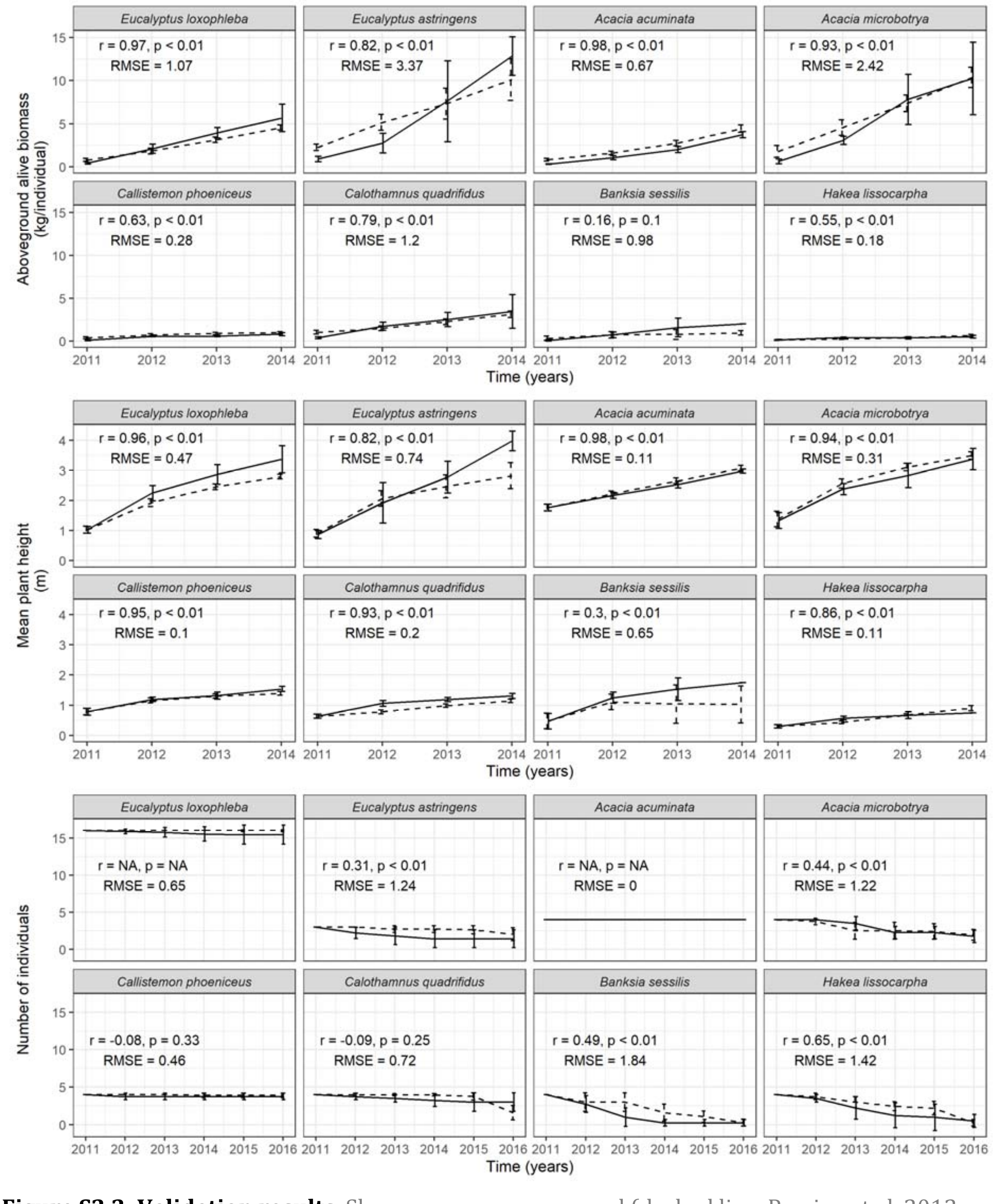


**Figure S2.2: Validation results.** Shown are mean measured (dashed line, Perring et al. 2012 and unpublished data) and mean simulated data (solid line) for aboveground alive biomass, plant height and population size over time and for the eight plant species planted in Ridgefield. Error bars show standard deviation over all repetitions for the assemblages plus in case for the simulation results over 10 model repetitions accounting for random plant height initialisation of neighbouring individuals to account for edge effects. Shown are Spearman's rank correlation coefficient *r*, corresponding p-value as well as the root mean square error *RMSE*. *NAs* for the correlation between measured and simulated number of individuals resulted when individual numbers over time were constant.

*References*

Bureau of Meteorology. 2019. http://www.bom.gov.au/. Accessed 11th March 2019.

Dentener, F., Drevet, J., Lamarque, J.F., Bey, I., Eickhout, B., Fiore, a. M., Hauglustaine, D., Horowitz, L.W., Krol, M., Kulshrestha, U.C., Lawrence, M., Galy-Lacaux, C., Rast, S., Shindell, D., Stevenson, D., Van Noije, T., Atherton, C., Bell, N., Bergman, D., Butler, T., Cofala, J., Collins, B., Doherty, R., Ellingsen, K., Galloway, J., Gauss, M., Montanaro, V., Müller, J.F., Pitari, G., Rodriguez, J., Sanderson, M., Solmon, F., Strahan, S., Schultz, M., Sudo, K., Szopa, S., Wild, O., 2006. Nitrogen and sulfur deposition on regional and global scales: A multimodel evaluation. Global Biogeochem. Cycles 20, n/a-n/a. https://doi.org/10.1029/2005GB002672

Harris, R.J., Standish, R.J., 2008. Ant dispersal and predation affects the availability of seeds for old-field recolonisation in Western Australia. J. R. Soc. West. Aust. 91, 301–311.

Hartig, F., Minunno, F., Paul, S., 2019. BayesianTools: General-Purpose MCMC and SMC Samplers and Tools for Bayesian Statistics. https://cran.r-project.org/package=BayesianTools.

Kattge, J., Díaz, S., Lavorel, S., Prentice, I.C., Leadley, P., Bönisch, G., Garnier, E., Westoby, M., Reich, P.B., Wright, I.J., Cornelissen, J.H.C., Violle, C., Harrison, S.P., Van BODEGOM, P.M., Reichstein, M., Enquist, B.J., Soudzilovskaia, N.A., Ackerly, D.D., Anand, M., Atkin, O., Bahn, M., Baker, T.R., Baldocchi, D., Bekker, R., Blanco, C.C., Blonder, B., Bond, W.J., Bradstock, R., Bunker, D.E., Casanoves, F., Cavender-Bares, J., Chambers, J.Q., Chapin Iii, F.S., Chave, J., Coomes, D., Cornwell, W.K., Craine, J.M., Dobrin, B.H., Duarte, L., Durka, W., Elser, J., Esser, G., Estiarte, M., Fagan, W.F., Fang, J., Fernández-Méndez, F., Fidelis, A., Finegan, B., Flores, O., Ford, H., Frank, D., Freschet, G.T., Fyllas, N.M., Gallagher, R. V., Green, W.A., Gutierrez, A.G., Hickler, T., Higgins, S.I., Hodgson, J.G., Jalili, A., Jansen, S., Joly, C.A., Kerkhoff, A.J., Kirkup, D., Kitajima, K., Kleyer, M., Klotz, S., Knops, J.M.H., Kramer, K., Kühn, I., Kurokawa, H., Laughlin, D., Lee, T.D., Leishman, M., Lens, F., Lenz, T., Lewis, S.L., Lloyd, J., Llusià, J., Louault, F., Ma, S., Mahecha, M.D., Manning, P., Massad, T., Medlyn, B.E., Messier, J., Moles, A.T., Müller, S.C., Nadrowski, K., Naeem, S., Niinemets, Ü., Nöllert, S., Nüske, A., Ogaya, R., Oleksyn, J., Onipchenko, V.G., Onoda, Y., Ordoñez, J., Overbeck, G., Ozinga, W.A., Patiño, S., Paula, S., Pausas, J.G., Peñuelas, J., Phillips, O.L., Pillar, V., Poorter, H., Poorter, L., Poschlod, P., Prinzing, A., Proulx, R., Rammig, A., Reinsch, S., Reu, B., Sack, L., Salgado-Negret, B., Sardans, J., Shiodera, S., Shipley, B., Siefert, A., Sosinski, E., Soussana, J.-F., Swaine, E., Swenson, N., Thompson, K., Thornton, P., Waldram, M., Weiher, E., White, M., White, S., Wright, S.J., Yguel, B., Zaehle, S., Zanne, A.E., Wirth, C., 2011. TRY - a global database of plant traits. Glob. Chang. Biol. 17, 2905–2935. https://doi.org/10.1111/j.1365-2486.2011.02451.x

Perring, M.P., Standish, R.J., Hulvey, K.B., Lach, L., Morald, T.K., Parsons, R., Didham, R.K., Hobbs, R.J., 2012. The Ridgefield Multiple Ecosystem Services Experiment: Can restoration of former agricultural land achieve multiple outcomes? Agric. Ecosyst. Environ. 163, 14–27. https://doi.org/10.1016/j.agee.2012.02.016

Rawls, W.J., Ahuja, L.R., Brakensiek, D.L., Shirmohammadi, A., 1992. Infiltration and soil water movement, in: Maidment, D.R. (Ed.), Handbook of Hydrology. McGraw-Hill Professional.

Rawls, W.J., Gimenez, D., Grossman, R., 1998. Use of soil texture, bulk density, and slope of the water retention curve to predict saturated hydraulic conductivity. Trans. Am. Soc. Agric. Eng. 41, 983–988. https://doi.org/10.13031/2013.17270

Schaphoff, S., von Bloh, W., Rammig, A., Thonicke, K., Biemans, H., Forkel, M., Gerten, D., Heinke, J., Jägermeyr, J., Knauer, J., Langerwisch, F., Lucht, W., Müller, C., Rolinski, S., Waha, K., 2017. LPJmL4 - a dynamic global vegetation model with managed land: Part I - Model description. Geosci. Model Dev. Discuss. 1–59. https://doi.org/10.5194/gmd-2017-145

Sitch, S., Prentice, I.C., Smith, B., Cramer, W., Kaplan, J.O., Lucht, W., Sykes, M., Thonicke, K., Venevsky, S., 2000. LPJ - A coupled model of vegetation dynamics and the terrestrial carbon cycle. Role Veg. Dyn. Control Atmos. CO2 Content, Ph.D. Thesis.

Sitch, S., Smith, B., Prentice, I.C., Arneth, a., Bondeau, a., Cramer, W., Kaplan, J.O., Levis, S., Lucht, W., Sykes, M.T., Thonicke, K., Venevsky, S., 2003. Evaluation of ecosystem dynamics, plant geography and terrestrial carbon cycling in the LPJ dynamic global vegetation model. Glob. Chang. Biol. 9, 161–185. https://doi.org/10.1046/j.1365-2486.2003.00569.x

Smith, B., Wärlind, D., Arneth, A., Hickler, T., Leadley, P., Siltberg, J., Zaehle, S., 2014. Implications of incorporating N cycling and N limitations on primary production in an individual-based dynamic vegetation model. Biogeosciences 11, 2027–2054. https://doi.org/10.5194/bg-11-2027-2014

Thomson, F.J., Moles, A.T., Auld, T.D., Kingsford, R.T., 2011. Seed dispersal distance is more strongly correlated with plant height than with seed mass. J. Ecol. 99, 1299–1307. https://doi.org/10.1111/j.1365-2745.2011.01867.x

### S3: Simulation experiments

#### S3.1: Processing of weather data from Pingelly

The weather data from Pingelly (Bureau of Meteorology, 2019, see main text *Climate Change Scenarios*) had to be corrected by filling data gaps and removing leap days.

Data gaps of up to two subsequent days were filled by the mean of the previous and the next day. For more than two missing days, we filled the gaps by taking the mean of the same days from the previous and the following year. If there was missing data in the previous or the following year, data gaps were only filled by the value of the year that has data available.

Leap days were deleted in order to allow for random weather year selection. In this case, precipitation events occurring on the leap day were added the next day. Precipitation events on the 28th of February were moved to the previous day to avoid doubling of these events when re-adding a leap day in the final weather input data set.

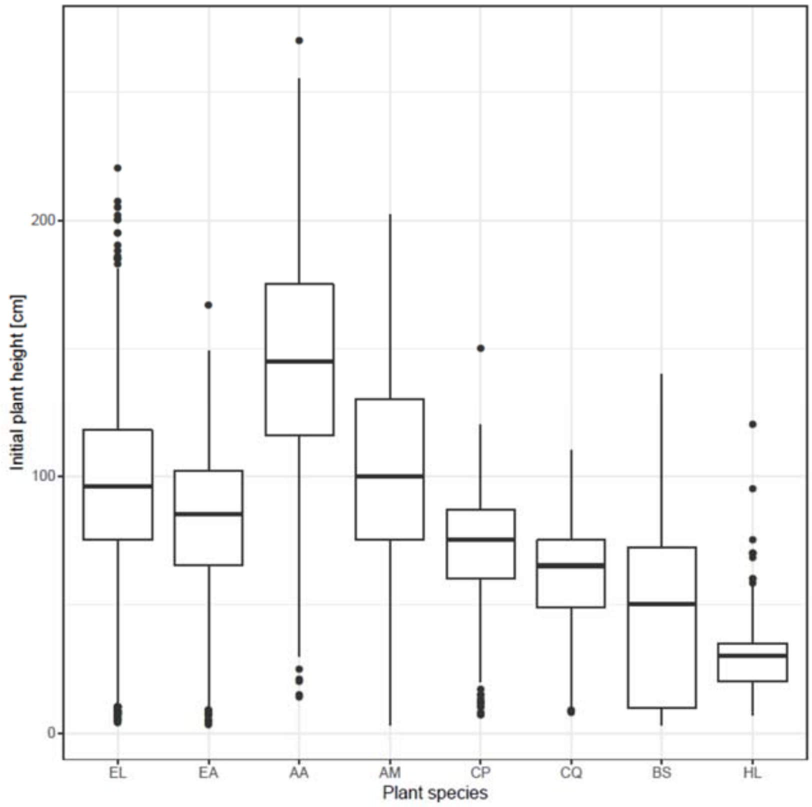


**Figure S3.1: Plant heights of the one-year old eight species in the Ridgefield experiment.** Data derived from Perring et al. (2012). EL: *Eucalyptus loxophleba,* EA: *E. astringens*, AA: *Acacia acuminata*, AM: *A. microbotrya*, CP: *Callistemon phoeniceus*, CQ: *Calothamnus quadrifidus*, BS: *Banksia sessilis,* HL: *Hakea lissocarpha*.

**Table S3.1: Median annual and seasonal climate trends.** Simulated data derived from Hope et al., (2015). Values provide the difference between 1995 (1986-2005) and 2090 (2080-2099) for SW Australia and RCP 8.5. Assumed atmospheric $CO_2$ of RCP 8.5 is derived from IPCC (2014).

| | **Annual** | **Winter (Jun to Aug)** | **Spring (Sep to Nov)** | **Summer (Dec to Feb)** | **Autumn (Mar to May)** |
|---|---|---|---|---|---|
| **Mean air temperature [°C]** | +3.36 | +3.02 | +3.50 | +3.54 | +3.39 |
| **Minimum air temperature [°C]** | +3.19 | +2.72 | +3.23 | +3.50 | +3.31 |
| **Maximum air temperature [°C]** | +3.64 | +3.46 | +4.02 | +3.63 | +3.45 |
| **Precipitation [%]** | -16.00 | -25.82 | -31.70 | -4.22 | -2.29 |
| **Solar radiation [%]** | +1.90 | +5.95 | +2.17 | -0.30 | -0.22 |
| **Atmospheric $CO_2$ [ppm]** | +450 | - | - | - | - |

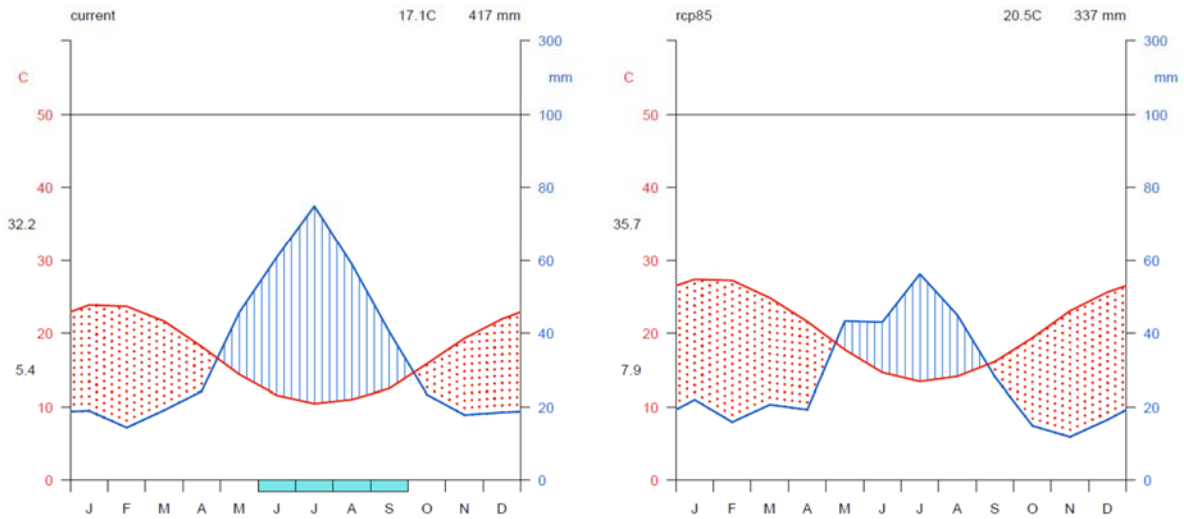


**Figure S3.2: Current (left) and future (right) climate diagrams for Pingelly.** Climate diagrams following the standard by Walter and Lieth (1969) created with R package by Guijarro (2019). Current climate (left) refers to the period 1990 and 2018 with mean annual precipitation of 417 mm as well mean annual temperature of 17.1 °C, future climate (right) refers to the same period but with projections of RCP 8.5 (see main text *Climate change scenarios*) with mean annual precipitation of 337 mm as well mean annual temperature of 20.5 °C . In each diagram monthly rainfall (blue) and temperature (red) is shown.

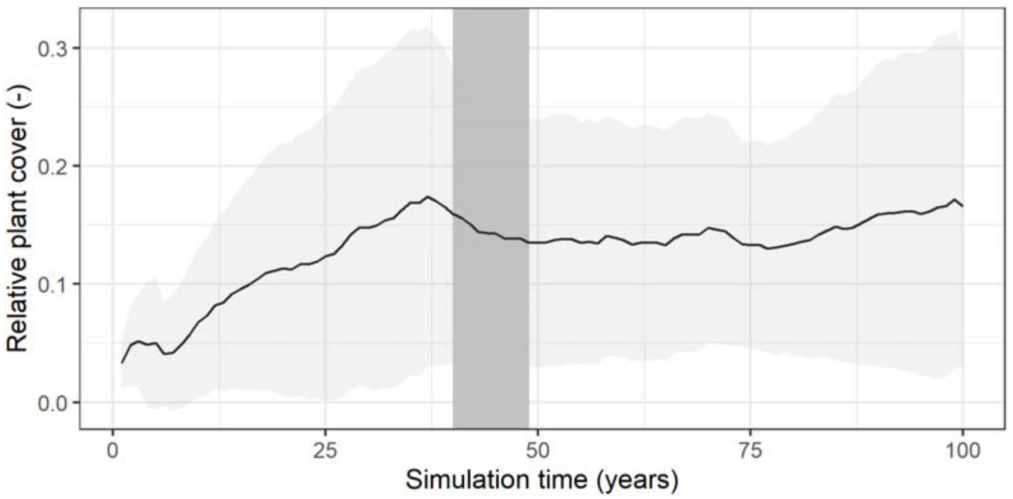


**Figure S3.3: Relative plant cover over time for the 8-species mixture under current climatic conditions.** Shown is mean and standard deviation over 10 model repetition. Rectangle between 40 and 49 simulation years highlights the time period during which simulation outcomes in this study where evaluated.

*References*

Bureau of Meteorology. 2019. http://www.bom.gov.au/. Accessed 11th March 2019.

Guijarro, J.A., 2019. climatol: Climate Tools (Series Homogenization and Derived Products). R package version 3.1.2. https://CRAN.R-project.org/package=climatol

Hope, P., Abbs, D., Bhend, J., Chiew, F., Church, J., Ekström, M., Kirono, D., Lenton, A., Lucas, C., McInnes, K., Moise, A., Monselesan, D., Mpelasoka, F., Timbal, B., Webb, L., Whetton, P., 2015. Southern and South-Western Flatlands Cluster Report, Climate Change in Australia Projections for Australia's Natural Resource Management Regions: Cluster Reports. https://doi.org/978-1-4863-0429-5

Perring, M.P., Standish, R.J., Hulvey, K.B., Lach, L., Morald, T.K., Parsons, R., Didham, R.K., Hobbs, R.J., 2012. The Ridgefield Multiple Ecosystem Services Experiment: Can restoration of former agricultural land achieve multiple outcomes? Agric. Ecosyst. Environ. 163, 14–27. https://doi.org/10.1016/j.agee.2012.02.016

Walter, H., Lieth, H., (1967). Klimadiagramm-Weltatlas. Gustav Fischer Verlag, Jena.

### S4: Supporting results

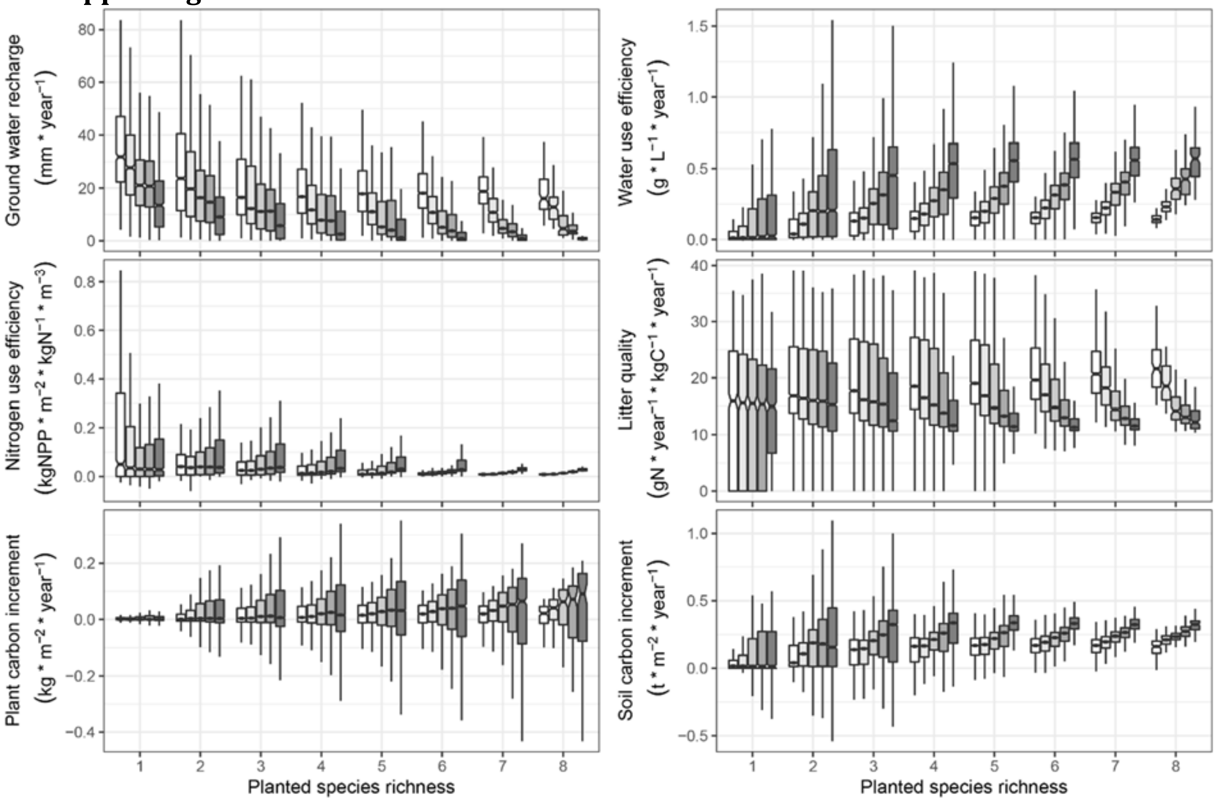


**Figure S4.1: Ecosystem functioning for each planted species richness under current (white boxplots) and different future climatic conditions (boxplots in different shades of grey in the order: RCP 26, 45, 60, and 85).** Shown is functioning for the last 10 simulated years and for 10 model repetitions as well as for 255 different plant communities which are unevenly distributed across the different planted species richness scenarios according to maximal possible combinations out of the pool of eight focal plant species. For better comparability among boxplots, single outliers are not shown.

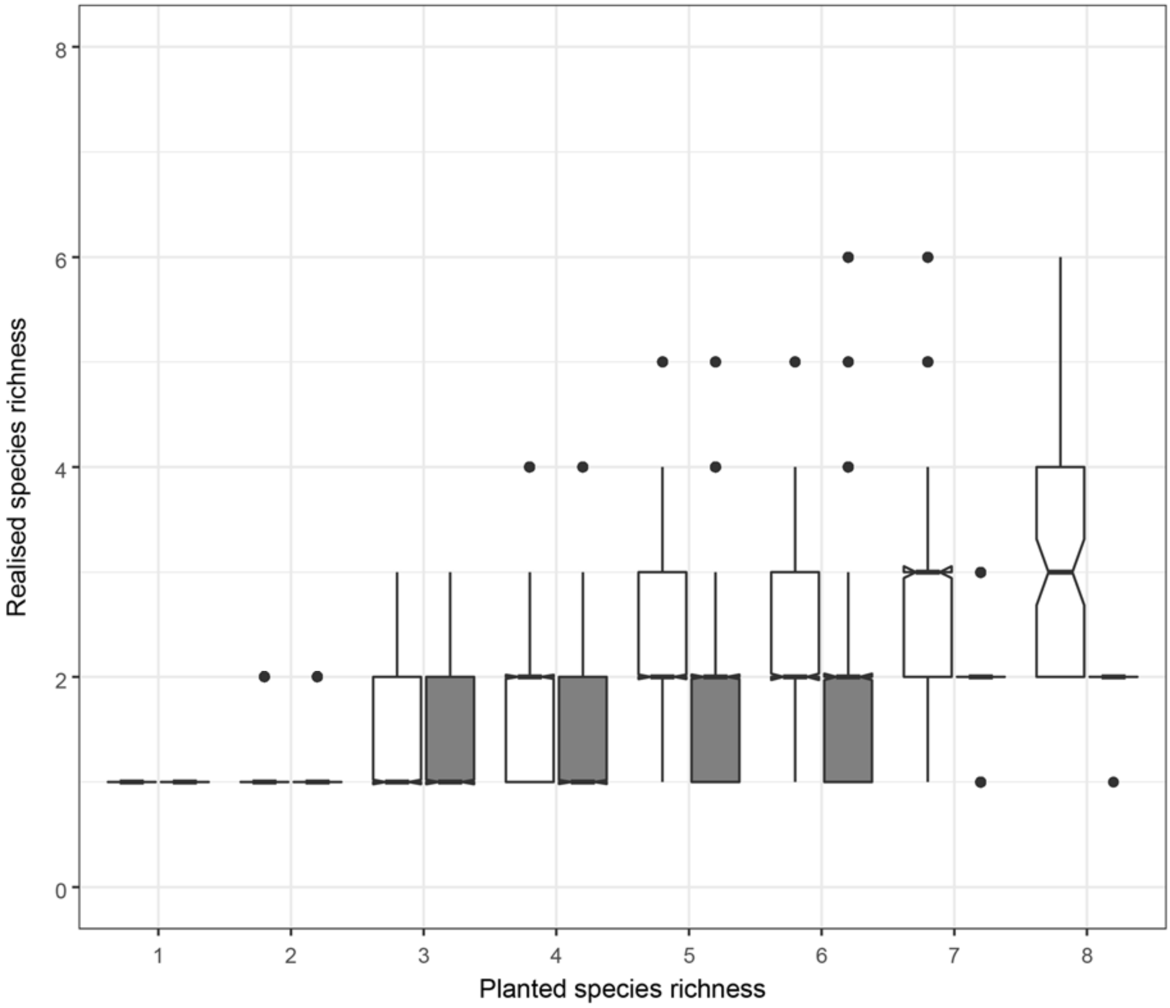


**Figure S4.2: Realised richness for each planted richness for current (white) and future conditions (grey).** Shown is realised richness over the last 10 simulated years and over 10 model repetitions.

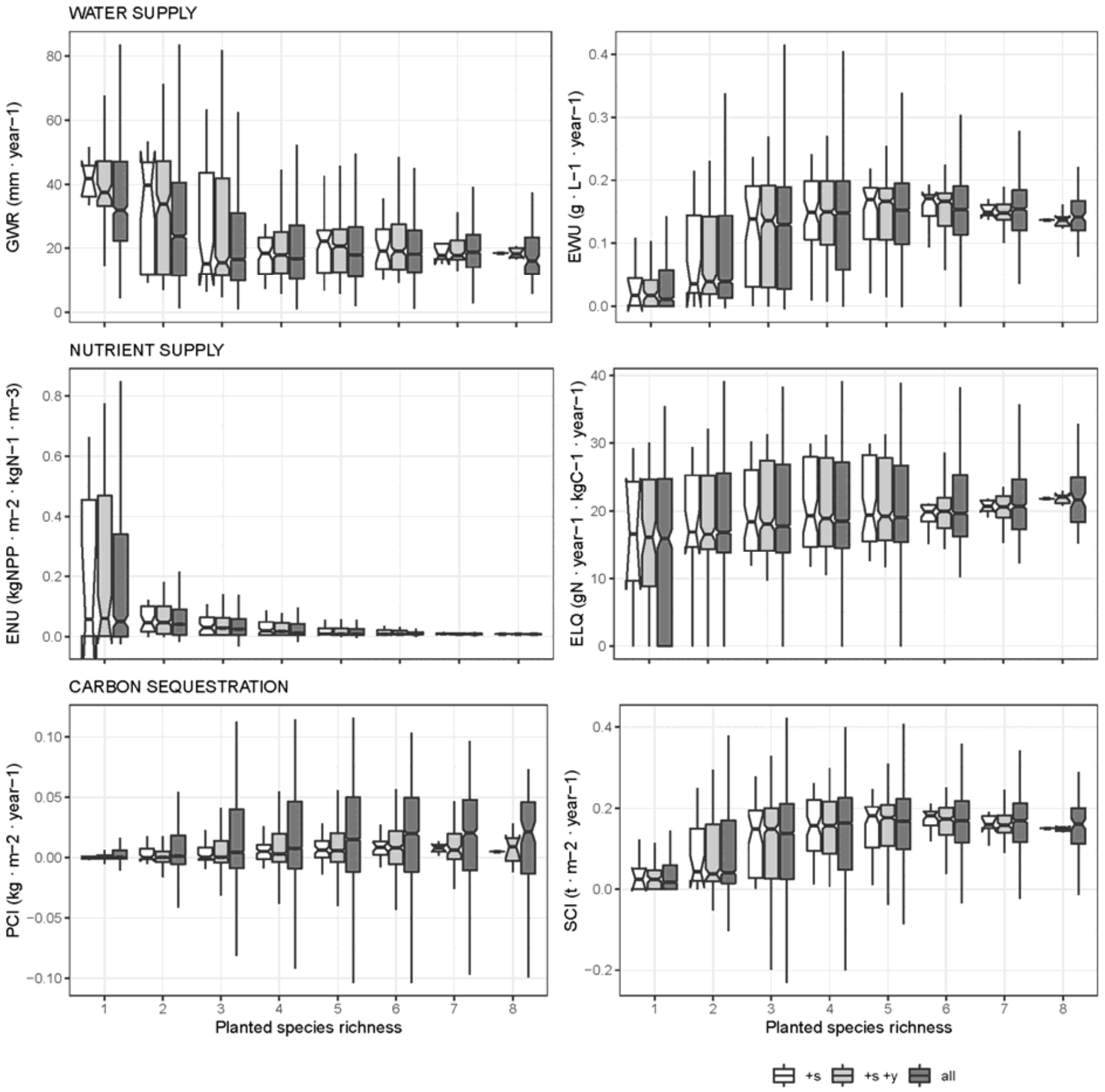


**Figure S4.3: Variability of ecosystem functioning under current conditions.** **+s**: mean functioning across years and model repetitions and thus represent variability of species compositions, **+s +y**: mean functioning over model repetitions and thus represent variability of species compositions across simulated years, **all**: functioning across species compositions, years and model repetitions (as shown in Fig. 2B in main text).

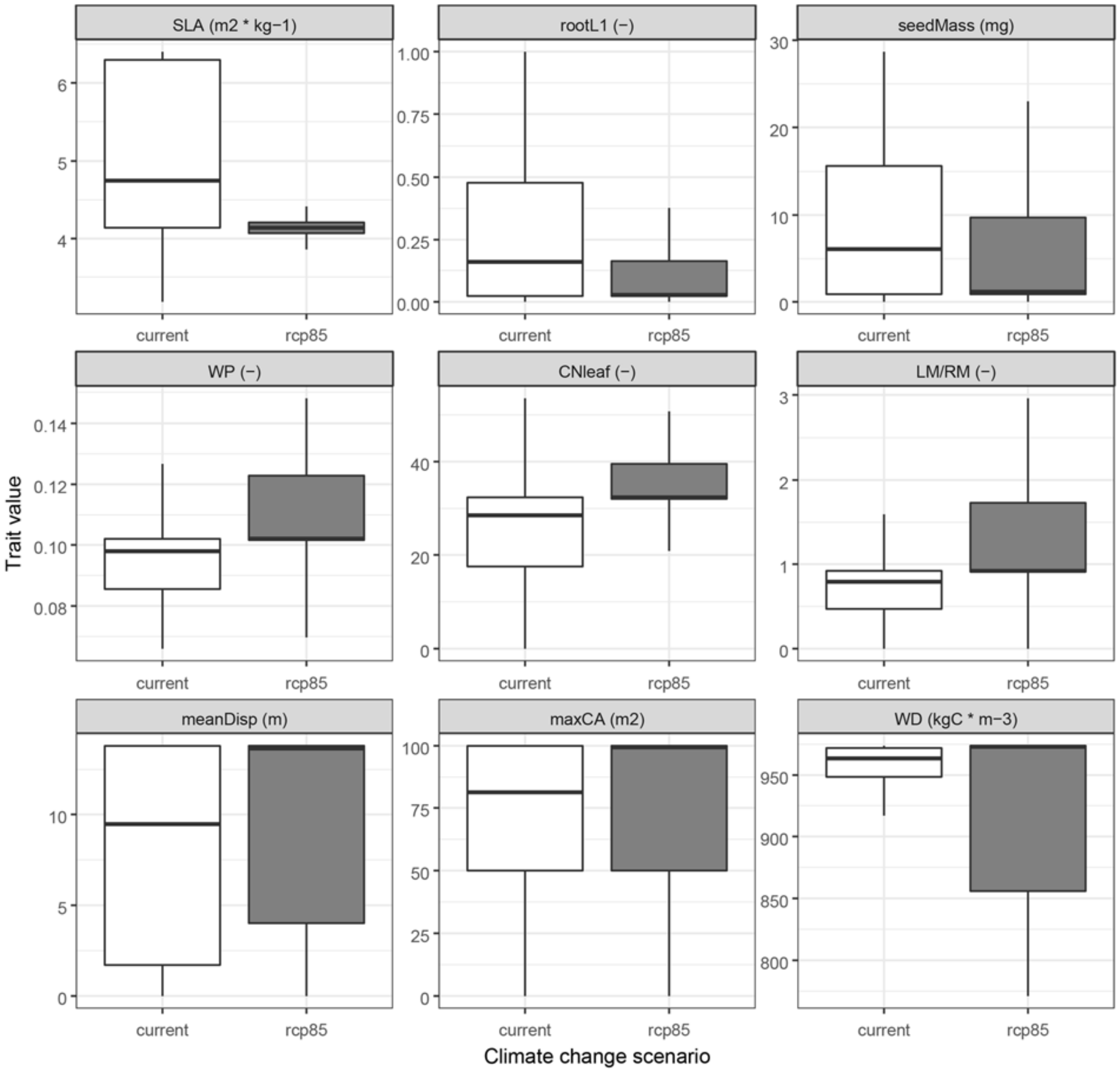


**Figure S4.4: CWM trait values across all planted species scenarios for current (white) and future conditions (grey).** Shown is CWM over the last 10 simulated years and over 10 model repetitions. Meaning of abbreviations can be found in Table 2 in the main text.

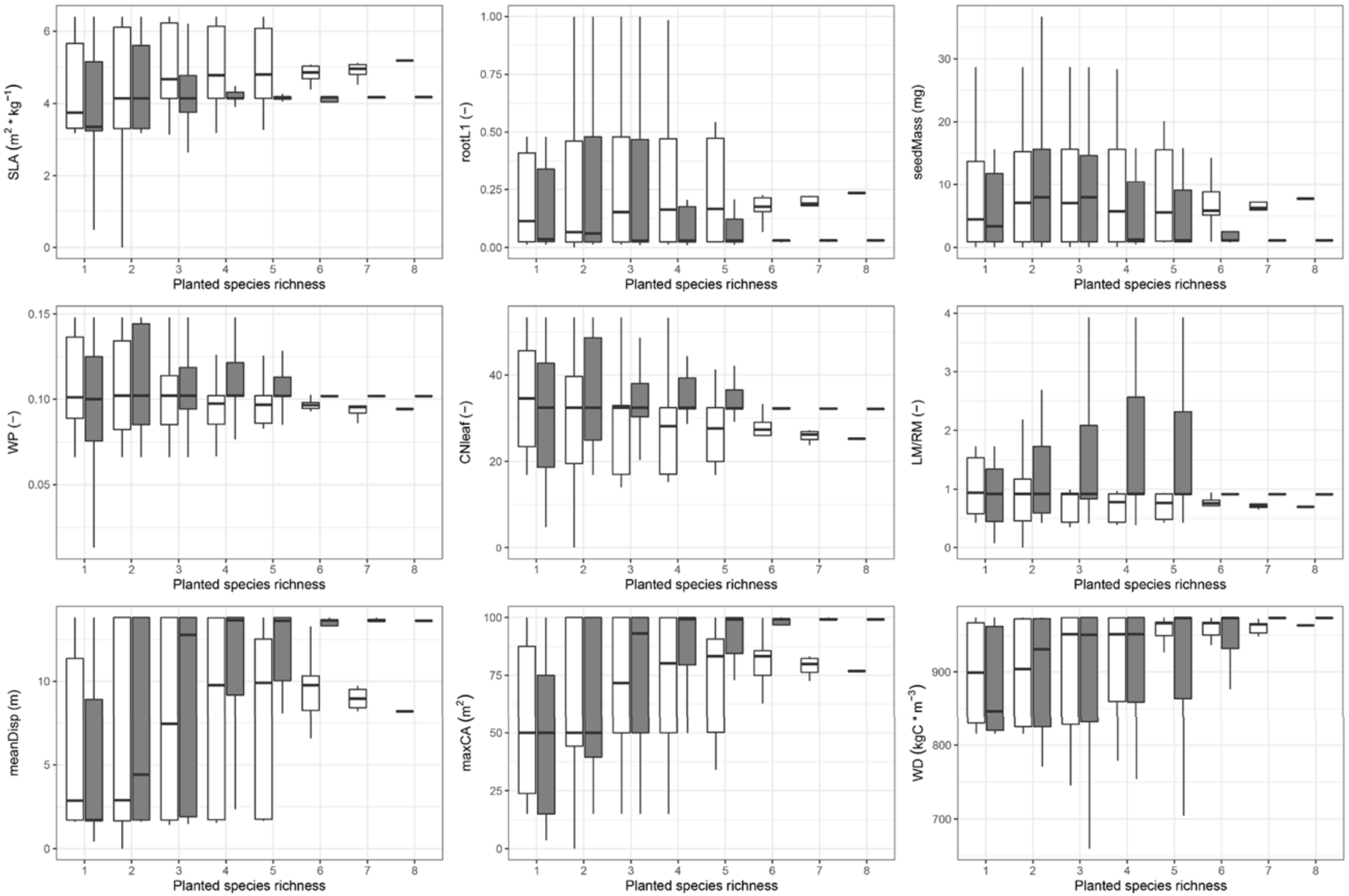


**Figure S4.5: CWM trait values for each planted richness for current (white) and future conditions (grey).** Shown is CWM over the last 10 simulated years and over 10 model repetitions. Meaning of abbreviations can be found in Table 2 in the main text.

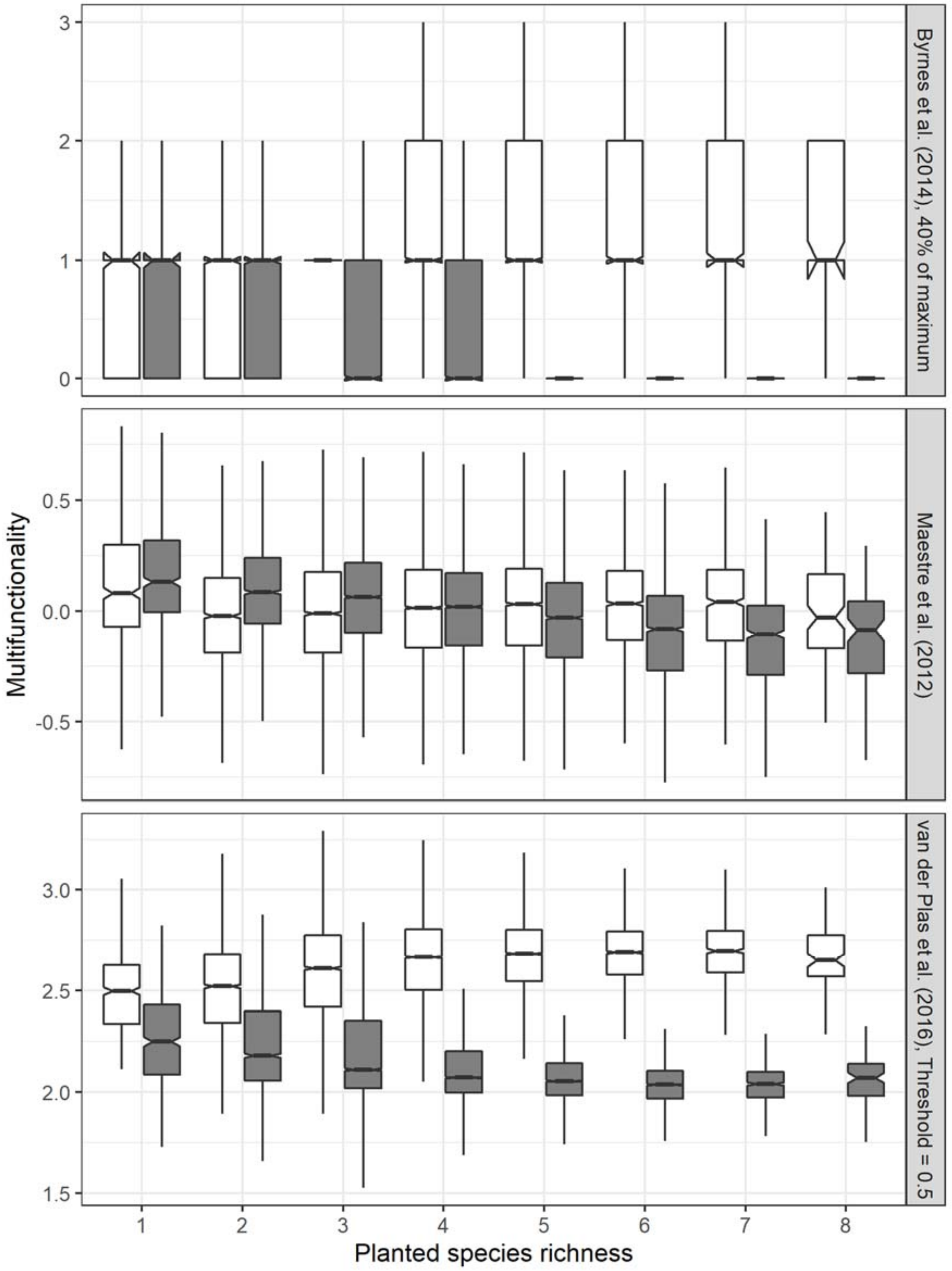


**Figure S4.6: Comparison of different multifunctionality indices across the different species richness and climate change scenarios (white: current, grey: future).** *Top panel:* Threshold approach by Byrnes et al (2014) counting functions with at least 40% of its maximum functioning. *Second panel*: Average over Z scores of the functions in a given climate scenario as suggested by Maestre et al. (2012), which accounts for the distance of each functioning to the mean of the same function. *Bottom panel*: Number of previously standardised functions between 0 and 1 based on the minimum and maximum value above an intermediate threshold of 0.5 in a given climate scenario as suggested by van der Plas et al. (2016) and used in this study. Comparison shows the approach by Byrnes et al. (2014) showed quantitatively similar trends as shown for van der Plas et al. (2016). In contrast, multifunctionality calculated after Maestre et al (2012) decreased with greater planted richness under current conditions.

*Reference list*

Byrnes, J.E.K., Gamfeldt, L., Isbell, F., Lefcheck, J.S., Griffin, J.N., Hector, A., Cardinale, B.J., Hooper, D.U., Dee, L.E., Emmett Duffy, J., 2014. Investigating the relationship between biodiversity and ecosystem multifunctionality: challenges and solutions. Methods Ecol. Evol. 5, 111–124. https://doi.org/10.1111/2041-210X.12143

Maestre, F.T., Quero, J.L., Gotelli, N.J., Escudero, A., Ochoa, V., Delgado-baquerizo, M., García-gómez, M., Bowker, M. a, Soliveres, S., Escolar, C., García-palacios, P., Berdugo, M., Valencia, E., Gozalo, B., Gallardo, A., Aguilera, L., Arredondo, T., Blones, J., Boeken, B., Bran, D., Conceição, A. a, Cabrera, O., 2012. Plant Species Richness and Ecosystems Multifunctionality in Global Drylands. Science 335, 2014–2017. https://doi.org/10.1126/science.1215442

van der Plas, F., Manning, P., Allan, E., Scherer-Lorenzen, M., Verheyen, K., Wirth, C., Zavala, M.A., Hector, A., Ampoorter, E., Baeten, L., Barbaro, L., Bauhus, J., Benavides, R., Benneter, A., Berthold, F., Bonal, D., Bouriaud, O., Bruelheide, H., Bussotti, F., Carnol, M., Castagneyrol, B., Charbonnier, Y., Coomes, D., Coppi, A., Bastias, C.C., Muhie Dawud, S., De Wandeler, H., Domisch, T., Finér, L., Gessler, A., Granier, A., Grossiord, C., Guyot, V., Hättenschwiler, S., Jactel, H., Jaroszewicz, B., Joly, F.-X., Jucker, T., Koricheva, J., Milligan, H., Müller, S., Muys, B., Nguyen, D., Pollastrini, M., Raulund-Rasmussen, K., Selvi, F., Stenlid, J., Valladares, F., Vesterdal, L., Zielínski, D., Fischer, M., 2016. Jack-of-all-trades effects drive biodiversity–ecosystem multifunctionality relationships in European forests. Nat. Commun. 7, 11109. https://doi.org/10.1038/ncomms11109
